# Group 15 Pre-emergent Herbicides Differentially Effect Plant Growth, Cuticular Wax Composition, and Fatty Acid Metabolism in Herbicide-Resistant and Herbicide-naïve Blackgrass

**DOI:** 10.1101/2024.09.20.614107

**Authors:** Hannah R. Blyth, Frederic Beaudoin, Richard P. Haslam, Barrie Hunt, Laurent Cornette, Dana R. MacGregor

**Affiliations:** Plant Sciences for the Bioeconomy, Rothamsted Research, Harpenden, AL5 2JQ, UK; Protecting Crops and the Environment, Rothamsted Research, Harpenden, AL5 2JQ, UK; Gowan Crop Protection Limited, Rothamsted Centre for Research and Enterprise, Harpenden, AL5 2JQ, UK

## Abstract

Despite their long history of effective use in agroecosystems, the precise molecular mechanisms of many pre-emergent herbicides are not fully understood. This study investigates the effects of three Group 15 pre-emergent herbicides (Flufenacet, S-ethyl dipropylthiocarbamate (EPTC), and tri-allate) on two well-characterized blackgrass (*Alopecurus myosuroides*) biotypes. Blackgrass is the predominant weed threatening winter wheat production in North-West Europe and the effective use of pre-emergent herbicides is important for preventing otherwise difficult-to-control blackgrass from establishing in agricultural fields. Using a sterile, agar-based system, we quantified the effects of multiple doses of flufenacet, EPTC, or tri-allate on plant physiology, including germination and early seedling growth, as well as the effects of a single dose on key biochemical pathways, specifically cuticular wax composition and fatty acid metabolism in biotypes exhibiting either non-target site herbicide resistance or complete sensitivity to all tested herbicides. Our data demonstrate that the three Group 15 pre-emergent herbicides alter cuticular wax composition and fatty acid profiles differently and that the resistant and herbicide-naïve biotypes show distinct responses to each herbicide. The GC-FID and GC-MS data from the different Group 15 pre-emergent herbicides are consistent with the observed differences in physiology and identify potential differences in how and where these herbicides act in these biotypes. Our findings provide novel molecular insights into the mechanisms of action of and tolerance to flufenacet, EPTC, or tri-allate in blackgrass.

## Introduction

Herbicides are chemicals that play an important role in controlling unwanted plant species (i.e. weeds) in agricultural ecosystems. They can be classified based on various factors, including their chemical structure, mode of action (MoA), timing of application, and target weed species. Herbicide classification is crucial for effective weed management, enabling growers and land managers to select herbicides with different sites of action for successive applications or mixtures, thereby improving control of target weeds while minimizing resistance selection pressures. Proper stewardship and integrated weed management practices are particularly important as the repeated use of the same herbicide or herbicides with a similar MoA can lead to the selection of herbicide-resistant weed populations (Hicks et al., 2018; Comont et al., 2019; Comont et al., 2020).

In 2020, the Herbicide Resistance Action Committee (HRAC) updated their MoA classification system, transitioning from letter to number MoA codes, adding missing herbicides, and updating the information to reflect new information (Liebl et al., 2020; Lerchl et al., 2024). This study focuses on three herbicides within the HRAC Group 15 herbicides, an α-oxyacetamide (flufenacet abbreviated FFT), and two thiocarbamates S-ethyl dipropylthiocarbamate (EPTC) and S-2,3,3-trichloroallyl di-isopropylthiocarbamate (tri-allate). Group 15 was created in 2020 (Liebl et al., 2020), reallocating herbicides previously characterised by a letter-based system as Group N, Z, and K3 (HRAC, 2010). Group 15 herbicides are also known as very-long-chain fatty acid (VLCFA) inhibitors, as they are suspected to target the biosynthesis of plant VLCFAs, i.e., fatty acids with more than 18 carbon atoms that are essential for the formation of waxes and important for developing the plant’s outer layer. For instance, data suggests EPTC inhibits wax deposition and cuticle formation by affecting the conversion of fatty acids into hydrocarbons and the elongation process (Gentner, 1966; Wilkinson and Hardcastle, 1970; Kolattukudy and Brown, 1974; Wilkinson and Smith, 1975; Gronwald, 1991). However, the biosynthetic enzyme or enzymes targeted by Group 15 herbicides remains largely unknown (Trenkamp et al., 2004). Moreover, HRAC’s official advice is that Group 15 herbicides may operate through a multi-site and/or multi-enzyme MoA, and there could be significant differences among herbicides within the group (Beffa et al., 2024).

To generate data that helps resolve this knowledge gap, we measured what effect(s) FFT, EPTC, and tri-allate have on plant physiology and fatty acid biosynthesis. Previous analyses comparing Group 15 herbicides have heterologously expressed Arabidopsis VLCFA elongases in *Saccharomyces cerevisiae* to compare the herbicides’ effects on VLCFA accumulation (Trenkamp et al., 2004). Their analysis showed that among the herbicides tested, flufenacet, allidochlor, and cafenstrole inhibited all VLCFA elongases tested and tri-allate, diphenamide, napropamide, and bensulide none (Trenkamp et al., 2004). In Trenkamp et al. (2004), EPTC was not tested and tri-allate was applied as a proto-herbicide, which requires additional metabolism *in planta* to have herbicidal activity (Schuphan and Casida, 1979); therefore conclusions drawn from this heterologous system may be complicated or incomplete. To increase our understanding of how these Group 15 pre-emergent herbicides work *in planta*, we assessed physiological and biochemical changes of blackgrass (*Alopecurus myosuroides*) exposed to three Group 15 herbicides under closed, sterile conditions. Flufenacet (Dücker et al., 2019; Dücker et al., 2020), EPTC (Fryer and Makepeace, 1978), and tri-allate (Allison, 2024) are all pre-emergent herbicides used for preventing blackgrass establishment in agricultural fields.

Blackgrass is an annual grass native to Europe and Asia, which has become a highly problematic weed in winter cereal crop production in Western Europe and China. Multiple-herbicide resistance is widespread in blackgrass (Hicks et al., 2018; Comont et al., 2020; Varah et al., 2020; Lan et al., 2022; Li et al., 2022; Qin et al., 2022) and resistance is highly heritable (Comont et al., 2022). Blackgrass adapts quickly to different management practices (Comont et al., 2019), and it exhibits tolerance to abiotic stress and these tolerances are correlated with increased herbicide resistance (Mohammad et al., 2022; Harrison et al., 2024). These traits alongside its ability to compete with crops for nutrients, water, and light means that blackgrass dramatically reduces crop yields and productivity when in the agroecosystem (Hicks et al., 2018; Varah et al., 2020; Zeller et al., 2021). There have been reports of multiple-herbicide resistant blackgrass biotypes in Germany and Sweden exhibiting resistance to Group 15 herbicides in 2007 and 2011 respectively (Heap, 2024). However, we focused on two well-characterised blackgrass biotypes that have been selected to exhibit archetype “Sensitive” or “Non-target Site Resistance (NTSR)” phenotypes. We are beginning to gain an understanding of the molecular mechanisms underpinning weedy traits in these populations (Mellado-Sánchez et al., 2020; Cai et al., 2023; Fu et al., 2023; Harrison et al., 2024), but their responses to Group 15 herbicides are unreported.

Reported instances of resistance to Group 15 herbicides are relatively low compared to other Groups – e.g. there are 10 reports of resistance to Group 15 herbicides compared to 175 reports for Group 2 (Heap, 2024). One of these is a report of wild oat that escapes tri-allate and has cross-resistance to difenzoquat (Group 8) (O’Donovan et al., 1994; Morrisson and Devine, 2005). Hierarchical clustering analysing cases of multiple-herbicide resistance showed HRAC Groups 15, Group 12 (inhibitors of Phytoene Desaturase) and Group 14 (inhibitors of Protoporphyrinogen Oxidase) were linked by their shared history of resistance (Hulme, 2022). Studies on blackgrass and *Lolium* sp. show that when used against different populations, FFT has varying efficacy, and resistance is linked to herbicide detoxification via glutathione S-transferase activity (Dücker *et al.,* 2019; Dücker *et al.,* 2019; Dücker *et al.,* 2020). The “Resistant” biotype we use here has NTSR that is conferred by enhanced herbicide detoxification via mechanisms including increased glutathione S-transferase activity (Cummins et al., 1997; Brazier et al., 2002; Mellado-Sánchez et al., 2020; Franco-Ortega et al., 2021).

Plant fatty acid synthesis has been reviewed extensively (see Li-Beisson et al. (2013) for details) and the production of VCLFA has been widely studied (reviewed in Batsale et al. (2021)). These data are summarised in Figure 1. Fatty acid synthesis begins in the chloroplast, and the main products are palmitic acid (C16:0; a 16-carbon fatty acid with no double bonds), stearic acid (C18:0), and oleic acid (C18:1). The first committed step in fatty acid synthesis is the formation of malonyl-CoA from acetyl-CoA and bicarbonate via acetyl-CoA carboxylase (ACCase), further synthesis of 16- or 18-carbon fatty acid is then performed by the fatty acid synthase complex (FAS). Plants contain both plastidic and cytosolic forms of ACCase. In grasses, the plastidic ACCase (a large multi domain protein) is homomeric, whereas in most dicots the plastidic form of ACCase is multimeric. The first activity of FAS includes condensing enzymes called 3-ketoacyl-ACP synthases (KAS) (Shimakata and Stumpf, 1982), and the successive addition of two-carbon units to the growing fatty acyl chain by two reductases and a dehydrase. The 3-ketoacyl-ACP is first reduced by a 3-ketoacyl-ACP reductase (KAR); 3-hydroxyacyl-ACP is then subjected to dehydration by the enzyme hydroxyacyl-ACP dehydratase (HAD), and then the resulting enoyl-ACP is finally reduced by the enzyme enoyl-ACP reductase (ENR) to form a saturated fatty acid. Some 16:0-ACP is released from FAS, but most molecules are elongated to 18:0 and are available for desaturation by the 18-ACP stearoyl-ACP desaturase (SAD). These newly synthesized fatty acids are subsequently incorporated into the glycerolipids of chloroplast membranes, or they are exported to the cytosol after termination of chain elongation by FATA and FATB acyl-ACP-thioesterases. Exported fatty acids are available for incorporation and modification into endoplasmic reticulum (ER) glycerolipids e.g., desaturation by fatty-acid desaturase-2 (FAD2), which produces linoleic acid (C18:2); and then fatty-acid desaturase (FAD3) to produce α-linolenic acid (C18:3).

**Figure 1:**
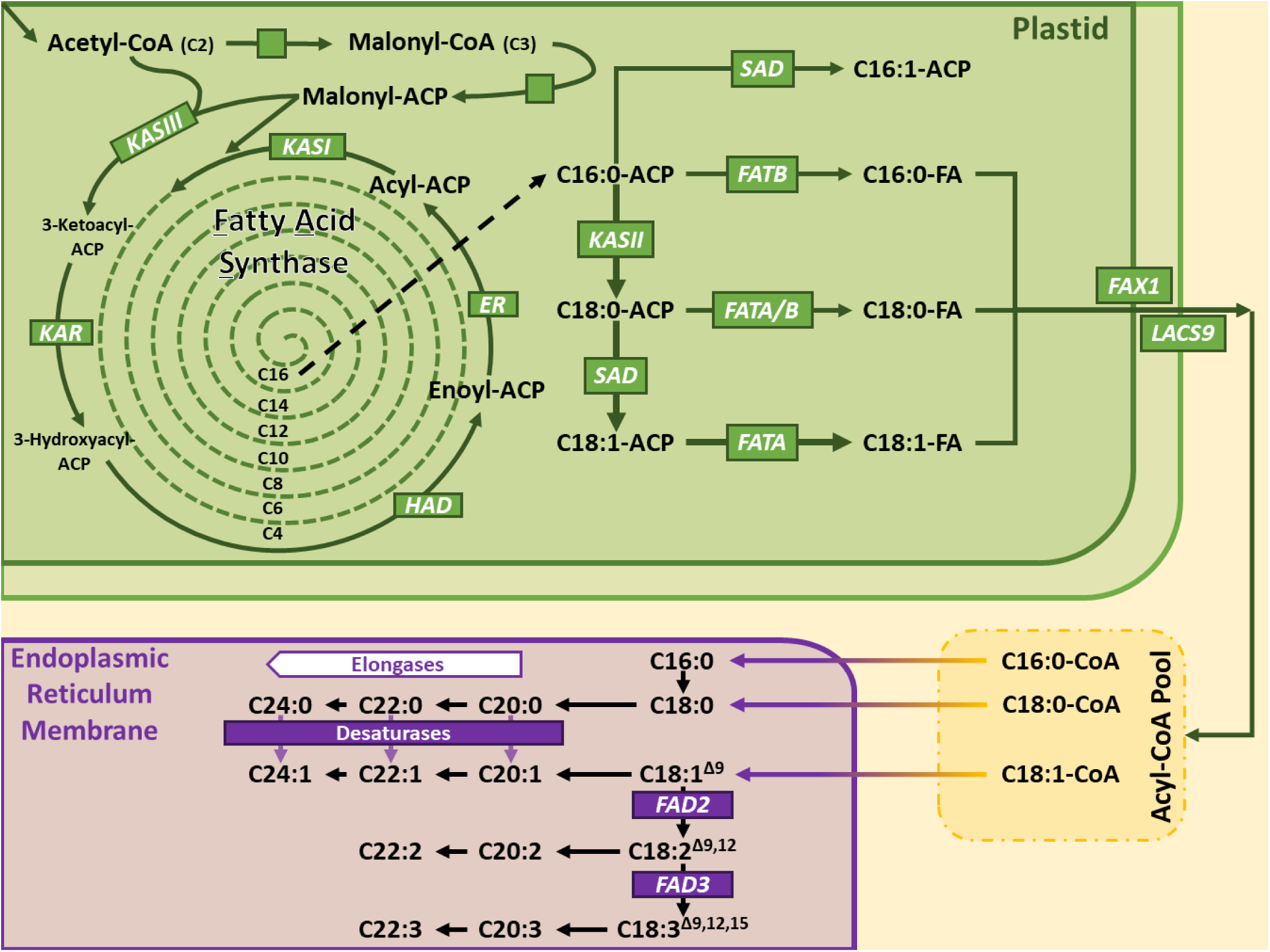
Schema representing the lipid biosynthetic pathway, indicating the putative impact of tri-allate treatment. This diagram illustrates the enzymatic steps involved in the elongation and desaturation of fatty acids, starting from C16:0-ACP (palmitoyl-ACP) in the plastid and ending with the production of very long-chain FAs with varying levels of desaturation in the ER. The formation of C16:0-ACP (palmitoyl-ACP) by the plastidial FAS complex is not shown. As described in the introduction this begins with the carboxylation of acetyl-CoA to form malonyl-CoA and is followed by an elongation cycle involving ACP-activated acyl chains. Each cycle consists in a condensation, a reduction, a dehydration, and a final reduction, progressively elongating the chain by 2 carbons until C16:0-ACP is formed. Represented on this diagram are: Ketoacyl-ACP Synthase III (KASIII), Ketoacyl-ACP Reductase (KAR), Hydroxyacyl-ACP Dehydrase (HAD), Enoyl-ACP Reductase (ER), Ketoacyl-ACP Synthase I (KASI), Ketoacyl-ACP Synthase II (KASII), Stearoyl-ACP Desaturase (SAD), Acyl-ACP Thioesterase B (FATB), Acyl-ACP Thioesterase A (FATA), Fatty Acid Export (FAX1), Long-Chain Acyl-CoA Synthetase (LACS9), Oleate Desaturase (FAD2), and Linoleate Desaturase (FAD3). Figure generated from figures and/or data at: http://aralip.plantbiology.msu.edu/data/tab_article.pdf or Li-Beisson et al. (2013).

Very long-chain fatty acid (VLCFA) biosynthesis begins when fatty acids exported to the cytosol, are activated as acyl-CoAs by long-chain acyl-CoA synthetases (LACS). LACSs are involved in numerous lipid processes, such as fatty acid transport and oxidation, and lipid synthesis (Shockey and Browse, 2011) and play a role in increased cuticular wax formation in response to changes in osmotic stress perceived by the roots (MacGregor et al., 2008). LACSs, therefore, serve fundamental roles in fatty acid metabolism. Long-chain acyl-CoAs represent the immediate precursors of VLCFAs and are further elongated through FAE (Fatty Acid Elongase) complexes localized in the endoplasmic reticulum membrane. Four core enzymes constitute the FAE complex and act in four sequential reactions to extend the acyl-chain by two carbon units (Haslam and Kunst, 2013). The first reaction consists of a condensation of the acyl-CoA substrate (n) with a malonyl-CoA catalysed by a condensing enzyme called 3-Keto-acyl-CoA Synthase (KCS) to give a 3-Keto-acyl-CoA (n+2). A 3-Keto-acyl-CoA Reductase (KCR) reduces the 3-keto group into an alcohol, to produce a 3-hydroxyacyl-CoA. Then a 3-Hydroxyacyl-CoA Dehydratase (HCD) converts this intermediate into an enoyl-CoA, which is reduced by an Enoyl-CoA Reductase (ECR) to form a saturated fatty acyl-CoA (n+2) product. This reaction cycle can be repeated to yield VLCFA with various chain lengths ranging from C20 up to C38 or more.

The seed specific KCS18 (FAE1) was the first FAE gene to be investigated and cloned in plants (Lemieux et al., 1990). Over recent years, genes coding for KCR, HCD, and ECR enzymes have been identified. KCR1 and PASTICCINO2 (PAS2) encode functional KCR and HCD enzymes, respectively, whilst CER10 encodes a functional ECR. Whilst the ECR subunit is often encoded by a single-copy gene, the condensing enzyme KCS is usually encoded by a large multi-copy gene family: 28 in maize (*Zea mays*) and as many as 58 in rapeseed (*Brassica napus*) (Batsale et al., 2023). As noted above, Trenkamp et al. (2004) tested 17 of the 21 elongases from Arabidopsis, including FAE1, KCS1, and KCS2, and showed that there is considerable differences in their specificity with respect to their elongation products and sensitivity to oxyacetamides, chloroacetanilides, and other compounds tested. In polyploid plants multiple-copy genes can encode the KCR and HCD components but with a much lower diversity. In Arabidopsis, PAS2 was characterised as the major HCD isoenzyme, but the PROTEIN TYROSIN PHOSPHATASE-LIKE (PTPLA) was later suggested as the HCD of specific FAE complexes expressed in vascular tissues that regulate endodermal VLCFA elongation by a yet-to-be-discovered mechanism (Morineau et al., 2016). Overall, the rather low genetic diversity found in KCR, HCD and ECR genes suggests that these three subunits generate large ranges of VLCFA chain lengths through broad substrate specificities, while KCS enzymes determine the chain-length substrate specificity of each elongation reaction (Kunst and Samuels, 2009; Batsale et al., 2023).

VLCFAs are important molecules that play crucial physiological and structural roles in plants. Membrane phospholipids such as Phosphatidyl-Choline (PC), Phosphatidyl-Ethanolamine (PE) and especially Phosphatidyl-Serine (PS) preferentially incorporate saturated C20, C22 and C24 VLCFAs, while sphingolipids accumulating in the outer leaflet of the plasma membrane (PM) are often enriched in α-hydroxylated saturated and monounsaturated C24 and C26 VLCFAs (Li-Beisson et al., 2013). Although most intracellular VLCFA have twenty-six or fewer carbon atoms (≤C26), plant cuticular VLCFAs and derivatives have chain lengths ranging from 26 to 34 carbon atoms, while leaf trichomes and pavement cells have been shown to produce VLCFA with up to 38 carbon atoms (Hegebarth et al., 2016). Plant species typically display variable amounts of free VLCFA in their cuticular waxes. Plant cuticular waxes are comprised of a range of chemical structures including fatty acids, aldehydes, primary and secondary alcohols, alkanes, ketones, and wax esters. Typically, wax components are synthesised by two distinct pathways (Bernard and Joubès, 2013): (1) the alcohol-forming (or reducing) pathway, in which WAX SYNTHASE/ACYL-COA: DIACYLGLYCEROL ACYLTRANSFERASE 1 (WSD1) catalyses the formation of wax esters using acyl-CoAs and primary alcohols as precursors; and (2) the conversion of very-long-chain acyl-CoAs into alkanes is catalysed by the ECERIFERUM1 (CER1)/ECERIFERUM3 (CER3)/CYTOCHROME B5 (CYTB5) complex in the alkane-forming pathway. Alkanes can be further oxidised to secondary alcohols and ketones by the CYP95A family of cytochrome P450 enzymes e.g., MIDCHAIN ALKANE HYDROXYLASE1 (MAH1). Wax components are then transported via trans-Golgi network (TGN)-trafficking pathways for export to the cuticle.

Our study revealed that these three Group 15 pre-emergent herbicides led to distinct physiological and metabolic phenotypes when applied to blackgrass grown in agar-based, sterile system. As predicted from field rate recommendations (Lainsbury, 2024), the amount of active ingredient required to elicit growth inhibition differed between the herbicides; FFT required the lowest doses, tri-allate the highest doses, while EPTC displayed intermediate efficacy. Only FFT efficiently impaired both shoot and root growth whereas both tri-allate and EPTC predominantly affected shoot growth. Moreover, Rothamsted and Peldon seeds responded differently to a given treatment. For both FFT and tri-allate, Rothamsted showed greater sensitivity than Peldon, as expected. Although the physiological effects of EPTC showed no major differences between control and treated plants, Rothamsted had smaller changes in wax and VLCFA composition, showing it was less affected by EPTC than Peldon. As expected for Group 15 herbicides, these herbicides reduced total leaf wax content. Of note, although there was little effect on root growth, tri-allate caused a decrease in total fatty acid content in shoots and roots that was not observed with EPTC or FFT. Our data highlight that each herbicide differentially affects physiology and lipid metabolism and that the two different blackgrass biotypes respond differently to a given treatment. Our data support the suggestion by Beffa et al. (2024) that these Group 15 herbicides operate through different MoAs and that the different biotypes respond differently to them, as we observe differences in efficacy, physiological impacts, biochemical changes, and biotype-specific responses between the Group 15 herbicides tested.

## Results

### Phenotypic characterisation of biotype responses to pre-emergent herbicides

Analysis of seeds sown on herbicide-impregnated agar supplemented with 0.85x Hoagland’s Solution showed that the two blackgrass biotypes responded differently to the separate pre-emergent herbicide and that the three pre-emergent herbicides inhibited shoot and/or root growth differently (Figure 2). FFT demonstrated the highest efficacy, requiring low doses to elicit a response (Figure 2, A-i & B-i and Supplementary Figure 1). Both shoot growth and root growth were impaired by FFT, although at equal doses, the relative inhibition of shoots was greater than of roots (Figure 2, A-i & B-i and Supplementary Figure 1). Rothamsted exhibited higher sensitivity to FFT than Peldon (Figure 2, A-i & B-i and Supplementary Figure 1). In contrast, reflecting field usage (Lainsbury, 2024), tri-allate required much higher herbicide doses to affect the blackgrass biotypes (Figure 2, A-ii & B-ii and Supplementary Figure 1). Rothamsted was more sensitive to tri-allate than Peldon (Figure 2, A-ii & B-ii and Supplementary Figure 1). Tri-allate’s impact on shoot growth was greater than on root growth (Figure 2, A-ii & B-ii and Supplementary Figure 1). EPTC showed intermediate efficacy, with effective concentrations falling between those of FFT and tri-allate (Figure 2, A-iii & B-iii and Supplementary Figure 1). The sensitivity of the two biotypes to EPTC was similar, with Peldon showing slightly higher sensitivity in most doses tested (Figure 2, A-iii & B-iii and Supplementary Figure 1). As with tri-allate, EPTC had a more pronounced effect on shoot growth than root growth (Figure 2, A-iii & B-iii and Supplementary Figure 1).

**Figure 2:**
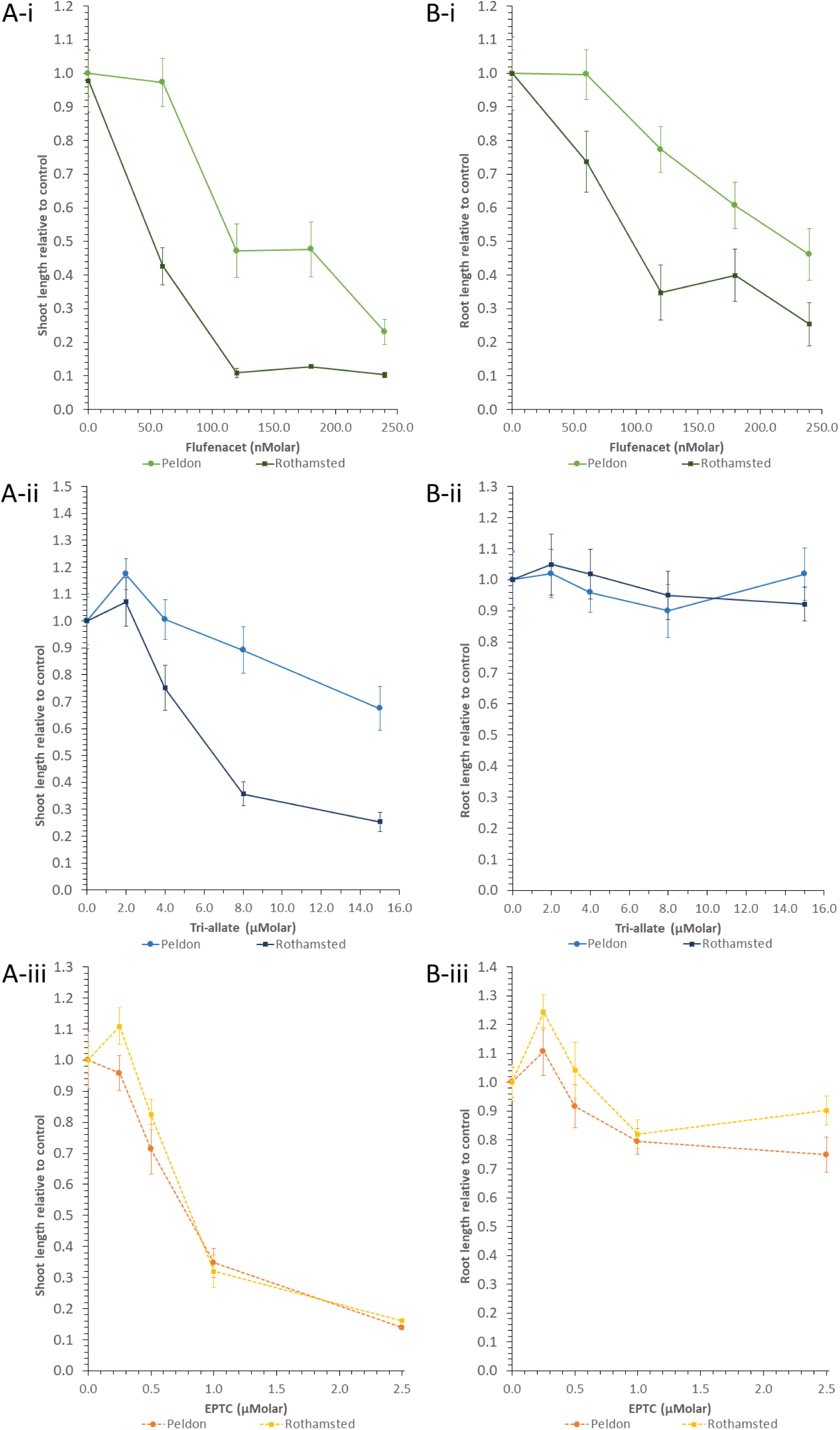
Example curves showing the different effects of the pre-emergent herbicides on the two blackgrass biotypes on (A) shoots and (B) roots. Showing average lengths relative to the biotype control ± standard error for (i) FFT; (ii) tri-allate; and (iii) EPTC.

Replicated experiments with different herbicide doses (Figure 2 and Supplementary Figure 1) allowed us to calculate an approximate ED40, equal to the dose that robustly altered plant growth without inducing mortality. The ED40 doses were estimated for the most sensitive biotype, and in the case of FFT, where the root growth phenotype had been affected by the herbicide, an additional ‘root’ ED40 was calculated. For EPTC and tri-allate, the shoot growth measurements were more sensitive indicators of herbicide effects than roots (Figures 2 & 3). These were 60 nM FFT shoots, 90 nM FFT roots, 600 nM for EPTC, and 6 μM for tri-allate (Figures 2 & 3). Figure 3 shows the phenotypic effects of the selected ED40 doses used to generate plant materials for wax and FAMES extractions. Indeed, the targeted response was achieved for the shoots in biotypes most sensitive to EPTC and tri-allate (Figure 3). In FFT, we successfully split the difference between Peldon and Rothamsted in both shoot and root tissues with the two doses (Figure 3).

**Figure 3:**
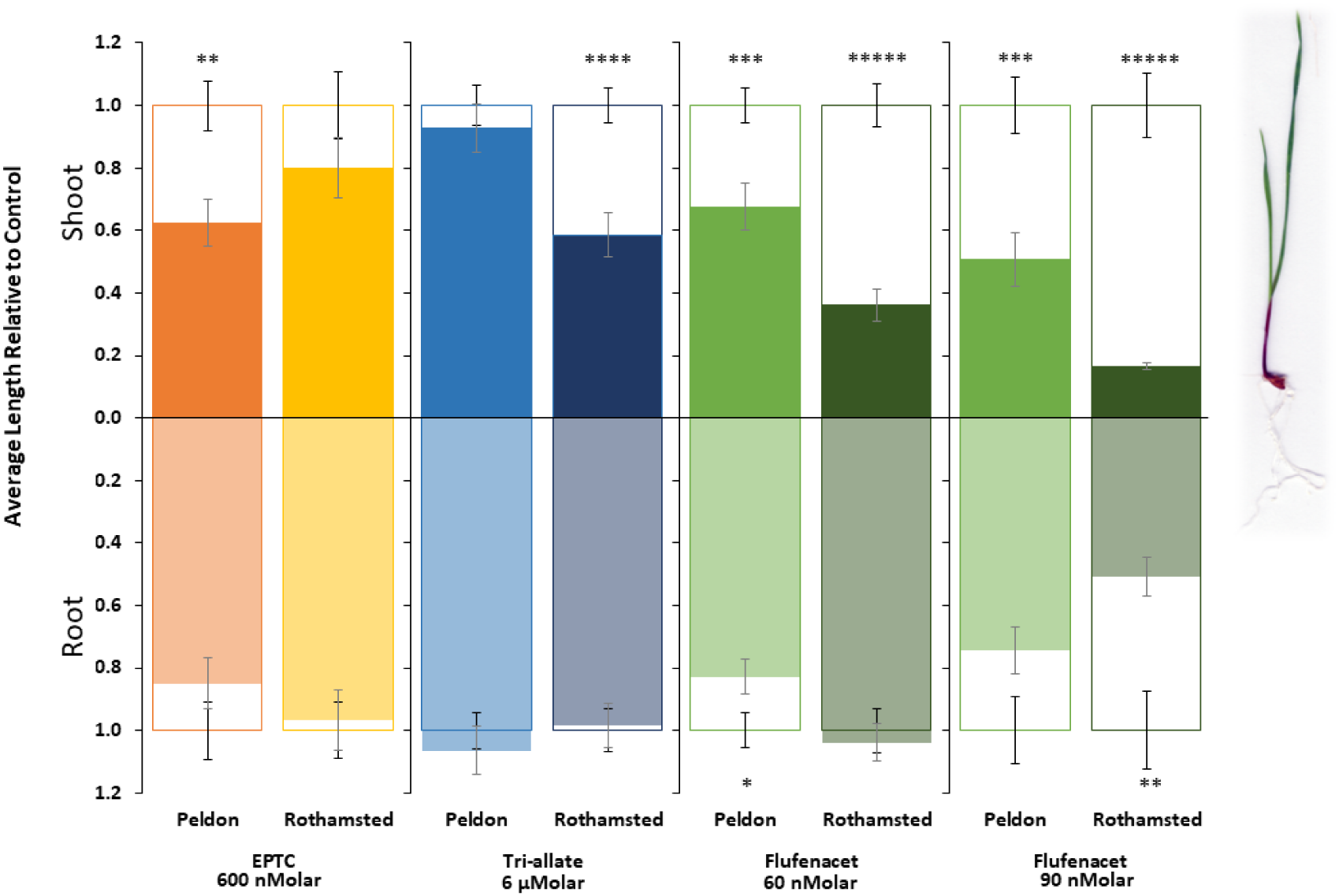
Bar plots showing the average length of shoots (up) and roots (down) relative to the untreated controls for each Blackgrass biotype and herbicide single-dose treatment. The outlines represent the control, while the fill represents the response to herbicide treatment at the selected respective dose: EPTC (600 nMolar), tri-allate (6 μM) and FFT (60 nM and 90 nMolar). Where the fill (herbicide) extends past the outline (control), the herbicide-treated samples are larger than the control. Error bars represent standard error. Significance codes results of paired t-tests comparing control to the herbicide-treated samples of the same biotype, * p ≤ 0.05 ** p ≤ 0.01 *** p ≤ 0.001 **** p ≤ 0.0001 ***** p ≤ 0.00001.

### Analysis of differences in total waxes with or without pre-emergent herbicides

In the closed, sterile, agar-based experimental conditions, the leaf surface wax load was 15.64 ng/mg for Peldon and 16.62 ng/mg for Rothamsted (average based on values for control treatments in Table 2). Our analyses revealed that blackgrass has an unusual leaf cuticular wax composition consisting of approximately 90 % very long chain fatty alcohols (VLC-FAOH), 6.5% fatty aldehyde and 3.5% alkanes (Table 1, average of controls). Hexacosanol (C26:0 FAOH) represents about 84% of total leaf wax (Supplementary Table 1, Figure 4). Only one molecular species of fatty aldehyde, hexacosanal (C26), was detected, and alkanes ranged from C27 to C35 (Table 1, Figure 4); although detected, values for C27 and C31 were close to the level of detection; therefore, they are not reported herein.

**Figure 4:**
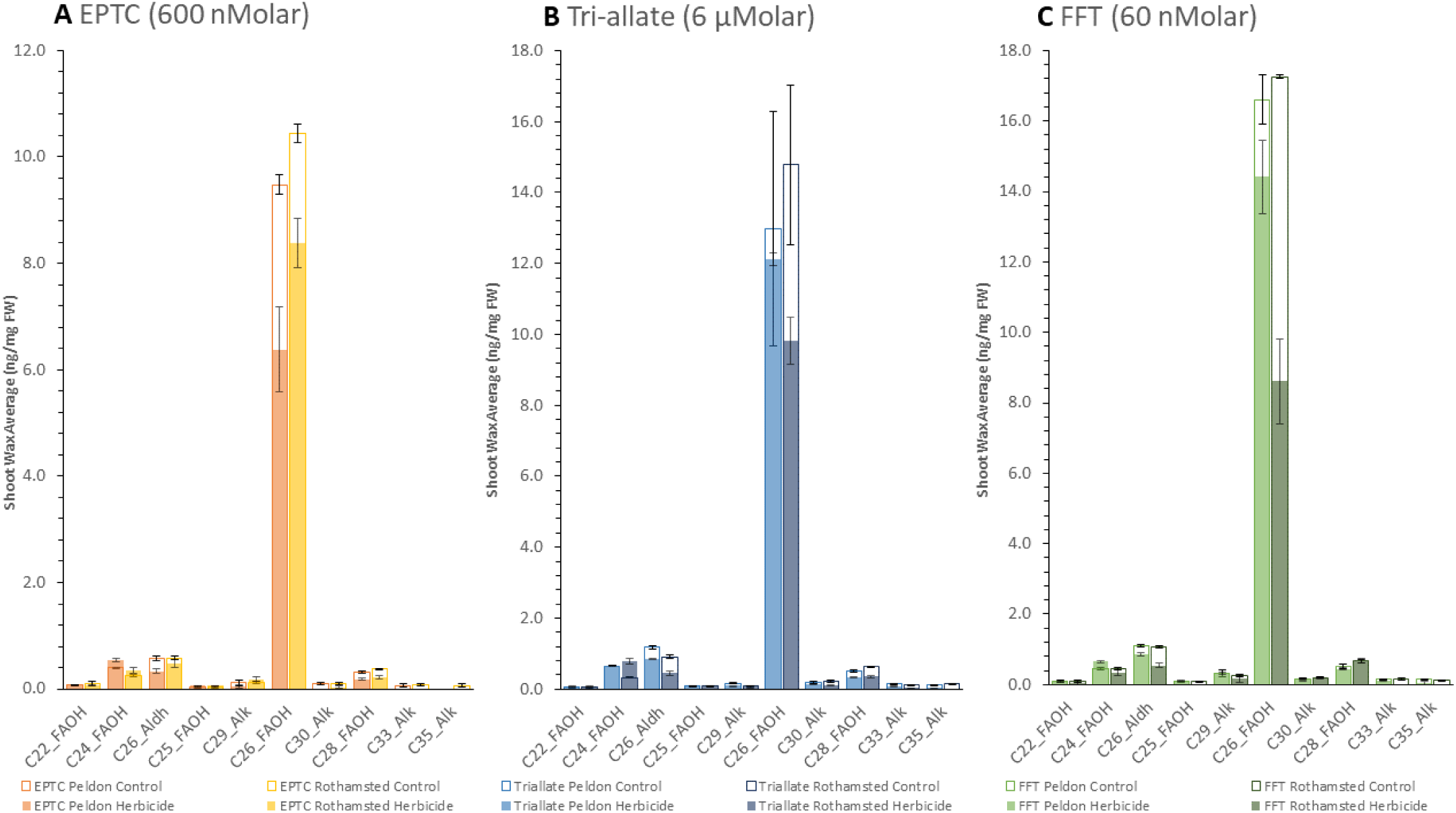
Rothamsted and Peldon shoot wax average composition ng/mg FW sample with and without herbicide treatment with (A) EPTC, (B) Tri-allate and (C) FFT. Analysis of data generated through GC-FID. The outlines represent the control, while the fill represents the response to herbicide treatment, n=3.

**Table 1:** Percentage composition of fatty alcohols (FAOH), aldehydes (ADH), and alkanes (ALK) in the total shoot wax content of two blackgrass biotypes.

| Herbicide | Biotype | Treatment | FAOH | ADH | ALK |
| --- | --- | --- | --- | --- | --- |
| EPTC | PELD | Control | 92.40 | 5.18 | 2.42 |
|  |  | Herbicide | 95.00 | 4.39 | 0.61 |
|  | Roth | Control | 92.12 | 4.81 | 3.08 |
|  |  | Herbicide | 92.95 | 4.83 | 2.22 |
| Tri-allate | PELD | Control | 87.77 | 8.06 | 4.16 |
|  |  | Herbicide | 91.87 | 5.83 | 2.31 |
|  | Roth | Control | 91.23 | 5.49 | 3.28 |
|  |  | Herbicide | 94.62 | 3.80 | 1.59 |
| FFT | PELD | Control | 90.21 | 5.68 | 4.11 |
|  |  | Herbicide | 91.48 | 4.97 | 3.55 |
|  | Roth | Control | 91.11 | 5.25 | 3.64 |
|  |  | Herbicide | 91.22 | 5.16 | 3.62 |

PELD, blackgrass biotype Pedon; Roth, blackgrass biotype Rothamsted.

**Table 2:** Average wax (ng/mg FW) species in blackgrass shoots, including sample weights and total wax amount.

| Herbicide | Biotype | Treatment | Average Sample Weight (mg) | C22-FAOH | C24-FAOH | C25-FAOH | C26-FAOH | C28-FAOH | C26-Aldh | C29-Alk | C30-Alk | C33-Alk | C35-Alk | Total Wax ng/mg FW |
| --- | --- | --- | --- | --- | --- | --- | --- | --- | --- | --- | --- | --- | --- | --- |
| EPTC | PELD | Control | 32.20 | 0.07 | 0.40 | 0.04 | 9.47 | 0.32 | 0.58 | 0.11 | 0.10 | 0.06 | 0.00 | 11.15 |
|  |  | Herbicide | 35.57 | 0.07 | 0.54 | 0.06 | 6.37 | 0.18 | 0.34 | 0.05 | 0.00 | 0.00 | 0.00 | 7.61 |
|  | Roth | Control | 46.77 | 0.09 | 0.25 | 0.03 | 10.45 | 0.38 | 0.58 | 0.13 | 0.10 | 0.09 | 0.06 | 12.15 |
|  |  | Herbicide | 45.17 | 0.07 | 0.35 | 0.06 | 8.38 | 0.22 | 0.48 | 0.18 | 0.03 | 0.00 | 0.00 | 9.76 |
| Tri-allate | PELD | Control | 34.67 | 0.06 | 0.65 | 0.10 | 12.98 | 0.51 | 1.18 | 0.17 | 0.19 | 0.14 | 0.10 | 16.07 |
|  |  | Herbicide | 49.90 | 0.07 | 0.66 | 0.08 | 12.13 | 0.32 | 0.84 | 0.09 | 0.14 | 0.09 | 0.02 | 14.43 |
|  | Roth | Control | 59.07 | 0.06 | 0.32 | 0.08 | 14.78 | 0.64 | 0.91 | 0.09 | 0.21 | 0.11 | 0.14 | 17.33 |
|  |  | Herbicide | 39.47 | 0.07 | 0.77 | 0.08 | 9.82 | 0.35 | 0.45 | 0.07 | 0.10 | 0.02 | 0.00 | 11.72 |
| FFT | PELD | Control | 31.33 | 0.10 | 0.46 | 0.09 | 16.61 | 0.52 | 1.12 | 0.33 | 0.17 | 0.16 | 0.14 | 19.69 |
|  |  | Herbicide | 24.23 | 0.09 | 0.65 | 0.10 | 14.41 | 0.45 | 0.85 | 0.32 | 0.13 | 0.12 | 0.04 | 17.16 |
|  | Roth | Control | 35.53 | 0.09 | 0.45 | 0.09 | 17.26 | 0.67 | 1.07 | 0.26 | 0.20 | 0.15 | 0.13 | 20.37 |
|  |  | Herbicide | 21.63 | 0.03 | 0.34 | 0.00 | 8.60 | 0.67 | 0.54 | 0.17 | 0.19 | 0.00 | 0.00 | 10.54 |

PELD, blackgrass biotype Pedon; Roth, blackgrass biotype Rothamsted.

The application of any of the three herbicides decreased the total amount of surface wax in both biotypes (Table 1, Figure 4). Using a C22 Alkane as an internal standard, we quantified the amount of each wax species identified in the ng/mg FW sample. Consistent with the morphological data in Figure 3, the change in Rothamsted’s surface wax load in response to EPTC was less than that of Peldon, with a 19.7 % vs. 31.7 % reduction in total surface wax, respectively (Table 2, Figure 4). Also matching the morphological effects, the opposite was observed with FFT and tri-allate. FFT and tri-allate induced 48.3 % and 32.4 % reductions in surface wax load in Rothamsted but only 12.9 % and 10.2 % reductions in Peldon, respectively (Table 2, Figure 4). These differences are also illustrated for all molecular species of surface waxes in Figure 4, with a closer view of lower abundance species in Supplementary Figure 2.

The Group 15 herbicides affected surface wax composition differently (Table 1). FFT did not affect the proportion of total fatty alcohols, aldehydes, and alkanes in Rothamsted and Peldon. In contrast, decreases in the proportion of aldehyde and alkane were observed in Peldon after treatment with EPTC and in both biotypes after treatment with Tri-allate. Interestingly, the three herbicides tested affected wax molecular species with carbon chain lengths of 26 and over, while C22 and C24 compounds were not affected (Figure 4, Table 2).

### Analysis of differences in FAMES with or without pre-emergent herbicides

We generated accurate, quantitative data measuring fatty acid content in ng per mg fresh weight from both shoot and root tissues using pentadecanoic acid (C15:0) as an internal standard, and response factors calculated for each individual fatty acid using the 37 FAMEs standard (SUPELCO). Analysis of blackgrass shoots and roots grown on sterile agar without herbicides shows a fatty acid composition characteristic of photosynthetic and non-photosynthetic tissues respectively (Supplementary tables 3.1 and 3.2, controls). In green leaf tissue, the profile is dominated by α-linolenic acid (ALA, C18:3Δ^9,12,15^ n3; ca. 60 %), which is very abundant in thylakoid membranes in the chloroplast, followed by linoleic acid (LA, C18:2Δ^9,12^ n6; ca. 19 %) and palmitic acid (C16:0; ca. 11 %) (Supplementary Table 3.1, controls). Very long chain fatty acids (VLCFA) with chain lengths ranging from 20 to 26 carbons were detected by GC-FID in shoots, but these only represent 4-5 % of total fatty acids. In root tissue, linoleic acid is the dominant fatty acid (40-42 %), followed by α-linoleic acid (19-26 %) and an increased proportion of palmitic acid (19-22 %) compared to shoots (Supplementary Table 3.2, controls). C20-C26 fatty acids are also slightly more abundant in root tissue (7-9 %) because they are incorporated in the suberin polymer found in the endodermis (Casparian strip) and periderm.

Adding the Group 15 pre-emergent herbicides into the closed agar-based system showed tri-allate had the strongest effect on fatty acid composition in both shoots and roots, primarily characterised by a dramatic decrease in polyunsaturated fatty acid (PUFA, i.e. LA and ALA) content (Figure 5, Table 3.1 and 3.2). This effect of tri-allate was more pronounced in shoots and stronger in Rothamsted than Peldon (Figure 5, Table 3.1 and 3.2). We saw no changes in the molecular composition of C16 and C18 FAs in shoots after treatment with EPTC and FFT (Figure 5, A-i and C-i, Table 3.1). Similar effects were observed in roots for EPTC and FFT, where no changes in the composition of C16 and C18 fatty acids were detected (Figure 5, A-ii and C-ii; Table 3.2). Minor decreases in total VLCFAs could be observed, and similar to the effect on surface wax content and composition, this was more pronounced with EPTC in Peldon in shoots and with FFT in Rothamsted in both shoots and roots (Supplementary Figure 3, A-i/ii and C-i/ii; Supplementary Table 3). Interestingly, in contrast with the effect observed on surface wax composition, these herbicides may also affect the levels of C20-C24 fatty acids in total lipids (Supplementary Figure 3). However, these variations are difficult to quantify accurately because of the very low level of VLCFAs in shoots. The effect of FFT was more evident with decreased levels of all saturated VLCFAs (C20-C26) in Rothamsted, while in Peldon roots, only species VLCFAs with chain lengths of C24 or greater were reduced in response to FFT (Supplementary Figure 3, C-ii).

**Table 3.1:** GC-FID fatty acid content ng/mg FW in blackgrass shoots with and without herbicide treatment.

| Herbicide | Biotype | Treatment | Average<br>Sample<br>Weight (mg) | C14:0 | C16:0 | C16:1<br>D9 | C18:0 | C18:1 | C18:2<br>n6 | C18:3<br>n3 | C20:0 | C20:1<br>n11 | C22:0 | C22:1 | C24:0 | C24:1 | C26:0 | Total<br>ng/mg<br>FW |
| --- | --- | --- | --- | --- | --- | --- | --- | --- | --- | --- | --- | --- | --- | --- | --- | --- | --- | --- |
| EPTC | PELD | Control | 39.57 | 0.59 | 39.33 | 5.86 | 3.77 | 6.62 | 65.13 | 204.19 | 1.40 | 0.50 | 3.36 | 0.00 | 2.04 | 5.32 | 2.25 | 342.07 |
|  |  | Herbicide | 52.30 | 0.45 | 35.43 | 5.54 | 4.94 | 3.50 | 59.70 | 195.06 | 1.11 | 0.44 | 3.72 | 0.00 | 1.67 | 1.90 | 1.95 | 316.12 |
|  | Roth | Control | 56.43 | 0.31 | 31.69 | 5.77 | 4.26 | 2.12 | 58.22 | 185.85 | 1.20 | 0.45 | 2.65 | 0.00 | 1.49 | 4.16 | 2.34 | 301.57 |
|  |  | Herbicide | 42.00 | 0.56 | 33.37 | 5.35 | 3.72 | 2.64 | 58.90 | 169.13 | 1.06 | 0.54 | 3.07 | 0.00 | 1.55 | 4.31 | 2.26 | 287.64 |
| Tri-allate | PELD | Control | 46.73 | 0.56 | 38.54 | 6.18 | 2.25 | 2.49 | 65.55 | 228.33 | 1.73 | 0.60 | 3.58 | 0.09 | 1.80 | 3.82 | 3.28 | 360.26 |
|  |  | Herbicide | 48.13 | 0.53 | 37.57 | 4.67 | 3.16 | 1.21 | 21.15 | 42.27 | 2.37 | 0.68 | 4.93 | 0.71 | 1.98 | 5.81 | 3.01 | 131.86 |
|  | Roth | Control | 42.83 | 0.84 | 37.30 | 6.20 | 12.52 | 2.87 | 64.74 | 195.97 | 2.19 | 0.62 | 4.01 | 0.35 | 2.47 | 8.18 | 3.12 | 341.79 |
|  |  | Herbicide | 55.53 | 0.35 | 29.99 | 3.75 | 2.19 | 1.34 | 11.31 | 12.71 | 1.08 | 0.37 | 3.91 | 0.78 | 1.96 | 7.97 | 1.85 | 81.20 |
| FFT | PELD | Control | 28.17 | 0.54 | 59.99 | 9.20 | 6.78 | 4.14 | 103.43 | 351.77 | 3.21 | 0.81 | 5.99 | 0.00 | 3.32 | 9.53 | 5.73 | 566.33 |
|  |  | Herbicide | 19.27 | 0.96 | 59.58 | 7.47 | 9.28 | 3.85 | 98.07 | 300.48 | 2.49 | 0.54 | 6.15 | 0.40 | 2.84 | 6.16 | 4.82 | 505.40 |
|  | Roth | Control | 34.73 | 0.70 | 48.86 | 7.99 | 8.97 | 3.47 | 83.17 | 262.86 | 2.35 | 0.66 | 5.45 | 0.00 | 2.21 | 8.25 | 4.08 | 441.07 |
|  |  | Herbicide | 17.67 | 0.24 | 47.31 | 6.77 | 5.16 | 3.09 | 81.52 | 246.15 | 1.54 | 0.00 | 4.27 | 0.00 | 2.22 | 4.67 | 3.10 | 407.97 |

PELD, blackgrass biotype Pedon; Roth, blackgrass biotype Rothamsted.

**Table 3.2:** GC-FID fatty acid content ng/mg FW in blackgrass roots with and without herbicide treatment.

| Herbicide | Biotype | Treatment | Average<br>Sample<br>Weight (mg) | C14:0 | C16:0 | C16:1<br>D9 | C18:0 | C18:1 | C18:2<br>n6 | C18:3<br>n3 | C20:0 | C20:1<br>n11 | C22:0 | C22:1 | C24:0 | C24:1 | C26:0 | Total<br>ng/mg<br>FW |
| --- | --- | --- | --- | --- | --- | --- | --- | --- | --- | --- | --- | --- | --- | --- | --- | --- | --- | --- |
| EPTC | PELD | Control | 32.97 | 0.06 | 12.17 | 0.44 | 1.57 | 4.05 | 26.22 | 16.89 | 0.18 | 0.00 | 1.46 | 0.36 | 0.91 | 0.27 | 0.09 | 64.67 |
|  |  | Herbicide | 51.50 | 0.09 | 10.57 | 0.38 | 0.76 | 3.89 | 22.48 | 14.32 | 0.14 | 0.02 | 1.38 | 0.65 | 1.04 | 0.27 | 0.18 | 56.17 |
|  | Roth | Control | 58.20 | 0.18 | 12.52 | 0.30 | 2.00 | 1.82 | 25.10 | 13.43 | 0.35 | 0.02 | 1.25 | 0.71 | 1.00 | 0.42 | 0.58 | 59.68 |
|  |  | Herbicide | 51.53 | 0.21 | 10.82 | 0.33 | 1.39 | 2.17 | 22.67 | 12.45 | 0.20 | 0.00 | 1.17 | 0.60 | 0.80 | 0.41 | 0.19 | 53.41 |
| Tri-allate | PELD | Control | 24.15 | 0.74 | 30.73 | 0.59 | 5.58 | 3.86 | 64.25 | 37.45 | 1.08 | 0.60 | 4.31 | 0.78 | 3.58 | 1.20 | 1.25 | 156.00 |
|  |  | Herbicide | 28.77 | 0.58 | 29.94 | 0.52 | 2.44 | 3.20 | 56.61 | 30.03 | 1.26 | 0.46 | 4.05 | 1.26 | 3.16 | 1.23 | 1.24 | 135.97 |
|  | Roth | Control | 23.27 | 0.70 | 43.13 | 0.96 | 3.59 | 2.96 | 78.90 | 51.45 | 2.16 | 0.69 | 4.40 | 1.65 | 4.06 | 1.69 | 1.75 | 198.09 |
|  |  | Herbicide | 28.30 | 0.51 | 27.00 | 0.70 | 1.45 | 2.53 | 45.65 | 19.83 | 0.95 | 0.71 | 3.23 | 1.55 | 1.92 | 0.98 | 0.79 | 107.78 |
| FFT | PELD | Control | 42.80 | 0.05 | 10.65 | 0.07 | 1.76 | 2.20 | 20.05 | 10.78 | 0.34 | 0.00 | 1.70 | 0.78 | 1.13 | 0.23 | 0.33 | 50.07 |
|  |  | Herbicide | 28.10 | 0.28 | 13.00 | 0.35 | 4.37 | 2.42 | 23.92 | 10.79 | 0.46 | 0.00 | 1.77 | 0.93 | 0.35 | 0.00 | 0.00 | 58.64 |
|  | Roth | Control | 41.67 | 0.06 | 14.57 | 0.15 | 2.25 | 2.09 | 28.04 | 12.57 | 0.44 | 0.00 | 1.98 | 1.10 | 1.48 | 0.54 | 0.43 | 65.70 |
|  |  | Herbicide | 22.17 | 0.25 | 11.36 | 0.07 | 1.84 | 1.50 | 27.51 | 10.88 | 0.17 | 0.00 | 1.06 | 0.69 | 0.00 | 0.00 | 0.00 | 55.33 |

PELD, blackgrass biotype Pedon; Roth, blackgrass biotype Rothamsted.

**Figure 5:**
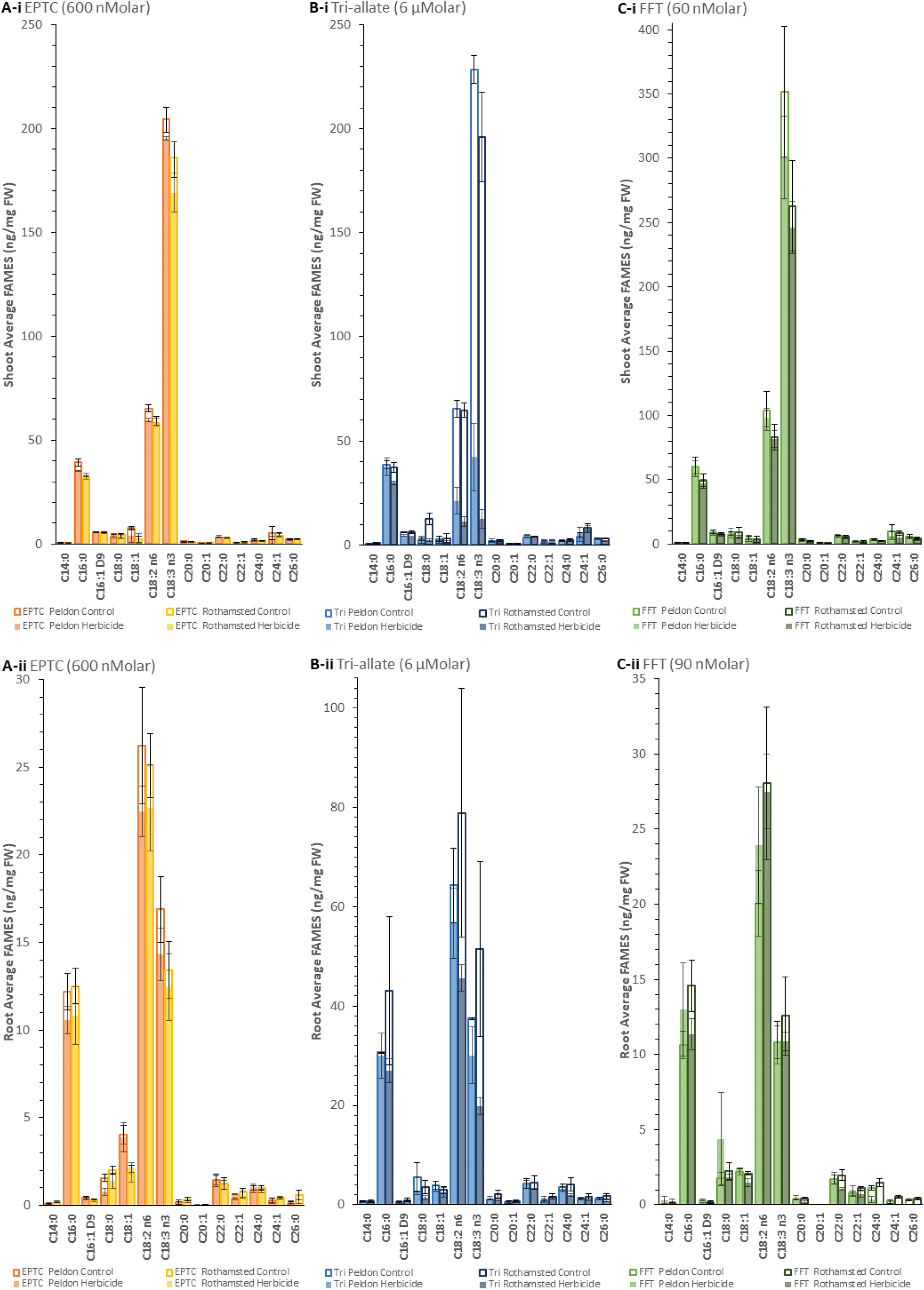
Quantified Differential Accumulation of FAMEs in Blackgrass (i) Shoot and (ii) Root Following Herbicide Treatment (A) EPTC, (B) Tri-allate (Tri) and (C) FFT. Analysed using GC-FID. Control samples not treated with herbicide are included for baseline comparison. Each data point represents the mean of three biological replicates, except for in (A-i) Peldon Herbicide treated and (B-ii) Tri-allate Control, where n=2, with error bars indicating a standard error.

Our quantitative analysis revealed a strong decrease in total fatty acid content in shoots and roots after treatment with tri-allate that was not observed with EPTC or FFT (Table 3.1, 3.2 and Supplementary Table 3). As described above, this reduction associated with tri-allate was more pronounced in shoots, with a 63.4 % decrease in Peldon and 76.2 % in Rothamsted, but also substantial in roots, with 12.8 % and 45.6 % decreases, respectively (Supplementary Table 3). This analysis also showed a sharp decrease in all unsaturated C18 fatty acids in shoots of both biotypes after treatment with tri-allate. In Peldon, there were 52.3 %, 67.7 % and 81.5 % reductions in C18:1Δ^9^ (oleic acid, OA), C18:2 (LA) and C18:3 (ALA) contents, respectively, and an even more substantial effect on Rothamsted with 55.0 %, 82.5 % and 93.5 % reduction in OA, LA and ALA content respectively (Figure 5, B-i; Table 3.1). Similar but less substantial decreases in OA, LA and ALA were also observed in roots (Table 3.2/Figure 5, B-ii), where the effect was greater in Rothamsted than Peldon.

Treatment with tri-allate also decreased Palmitoleic acid (PO, C16:1Δ^9^) content in both Peldon (24.5 % shoots and 11.8 % root) and Rothamsted (39.4 % shoots and 26.8 % roots) (Supplemental Figure 4). Inspection of palmitic acid (PA, C16:0) and stearic acid (SA, C18:0) content revealed that tri-allate affects the biotypes differently. In Peldon, the content of these two saturated fatty acids is not affected by tri-allate treatment in either shoots or roots, while in Rothamsted, PA and SA content are reduced in both shoots and roots (Figure 5, Table 3.1 and 3.2).

Analysis of VLCFA content and composition shows marked differences between biotypes and herbicides. At the dose used in this study, EPTC had minimal effect on VLCFA content in shoots and no effect on the root (Supplementary Figure 3, A-i and A-ii, Supplementary Table 3). FFT treatment decreased all saturated and mono-unsaturated VLCFAs in Rothamsted, but this effect was greater in the root, as discussed above. In Peldon, this reduction in VLCFA content only affected species of chain lengths of C24 or longer (C24+) in roots (Supplementary Figure 3, C-ii). Like FFT, tri-allate had very little effect on VLCFA content in shoots but affected all VLCFAs in Rothamsted roots (Supplementary Figure 3, B-i and B-ii).

## Discussion

Blackgrass is well known for its rapidly-evolved, widespread herbicide resistance. Extensive human-mediated gene flow between field populations (Dixon et al., 2020) has moved highly heritable resistances (Comont et al., 2022) conferred by NTSR mechanisms that are both shared and population-unique and encoded for on the chromosomes (Cai et al., 2023) and extra-chromosomal circular DNA (Fu et al., 2023) extensively around the UK (Comont et al., 2020) without correlated fitness penalties (Comont et al., 2022). Although the allele variants in herbicide targets are known (Comont et al., 2020), and we are beginning to derive a mechanistic understanding of NTSR mechanisms (Mellado-Sánchez et al., 2020; Cai et al., 2023; Fu et al., 2023; Lowe et al., 2024), there are still many questions about precisely how herbicides, particularly pre-emergent herbicides, effectively control plant growth.

We observed that without herbicide application, the cuticular wax composition of blackgrass is very unusual and different from most plants analysed to date, including Arabidopsis or mono- and dicotyledonous crops such as maize, rice, tomato, or rapeseed (Lee and Suh, 2015). A similar wax composition has only been reported recently for Taiwan oil millet, where the predominant wax component is a C28 primary alcohol in the leaf blade and a C28-free fatty acid in the leaf sheath (Anggarani et al., 2024).

Application of Group 15 pre-emergent herbicides, which are used to control blackgrass in agriculture and are predicted to alter VLCFA biosynthesis, led to changes in physiology and biochemistry in two well-characterised biotypes of blackgrass. The differences in herbicide doses required to induce these changes in our agar-based system (e.g. Figure 2) reflect the differences in recommended field rate use for these herbicides; e.g. recommended rates for field use of tri-allate are approximately 10X that of flufenacet (2250g tri-allate/ha compared to 240g flufenacet/ha (Lainsbury, 2024). The herbicide-naïve biotype used here (“Rothamsted”) originates from the Broadbalk long-term field experiment (Moss et al., 2004) and represents a population which has never been exposed to any herbicide. It is, therefore, particularly interesting that the growth of Rothamsted was less affected by the inclusion of EPTC in the agar (Figure 2Aiii) than was Peldon (Figure 2Biii), as Peldon is well-characterised for exhibiting strong NTSR to multiple herbicides across different groups and MoAs (Moss, 1990; Cummins et al., 1997; Hall et al., 1997; Mellado-Sánchez et al., 2020; Franco-Ortega et al., 2021; Comont et al., 2022; Cai et al., 2023; Fu et al., 2023). These differences in sensitivity to EPTC were correlated with differential changes in wax and VLCFA composition between the biotypes where the amount of change in Rothamsted was less than that in Peldon (Table 2, Figure 4) and the minor decreases in VLCFAs in shoots were more pronounced in Peldon than Rothamsted (Figure 5, A-i and C-i; Table 3.1). Our data are consistent with observations that EPTC inhibited wax deposition and cuticle formation by affecting the conversion of fatty acids into hydrocarbons and the elongation process (Gentner, 1966; Wilkinson and Hardcastle, 1970; Kolattukudy and Brown, 1974; Wilkinson and Smith, 1975; Gronwald, 1991). Moreover, the surprising result of Rothamsted being less affected than Peldon reiterates previous conclusions that weed biotypes should each be considered as individual populations (Hamidzadeh Moghadam et al., 2021; Hamidzadeh Moghadam et al., 2023), which may not respond in precisely predictable ways to control methods. However, because at the dose used in this study, EPTC had such a small effect on VLCFA content in shoots and no effect on the root (Figure 5, A-i and A-ii and Supplementary Table 3) to pinpoint where in the pathway EPTC is targeting and how targeting might differ between biotypes would require further investigation using additional herbicide doses.

Unlike EPTC, FFT and tri-allate more efficiently inhibited Rothamsted’s growth compared to Peldon (Figures 2 and 3). Like EPTC, tri-allate only altered shoot growth, while FFT repressed both root and shoots (Figures 2 and 3). FFT treatment decreased saturated and mono-unsaturated VLCFAs in Rothamsted, and this effect was greater than in the root (Figure 5). Figure 3 C-i shows that C20: and C20:1 are also affected by FFT in Peldon shoots while specifically C22:0/1 are unaffected. In Peldon, this reduction in VLCFA content was also generally observed in shoots but in roots only species of chain lengths of C24 or longer (C24+) are affected (Figure 5, C-i and C-ii). These Rothamsted data are consistent with previous data that showed FFT inhibits the activity of all tested Arabidopsis VLCFA elongases in a heterologous system (Trenkamp et al., 2004) whereas the Peldon data suggest it exhibits partial resistance which allows for C22 formation in the shoots and C20/C22 in the roots.

Like FFT, tri-allate had a small effect on VLCFA content in shoots and even though tri-allate did not affect root length, an effect on VLCFA content in Rothamsted roots (Figure 5, B-i and B-ii). Despite the small effect of tri-allate on VLCFA content in shoots, C26 was clearly decreased in both Rothamsted and Peldon shoots (Supplemental Figure 4 B-i), which is consistent with the observed decrease in C26+ wax species (Figure 4). Unexpectedly, the strongest effect of tri-allate on fatty acids content and composition in both biotypes was the strong reduction in C18 desaturated FAs.

These initially suggested tri-allate affects fatty acid desaturation (Figure 1). Treatment with tri-allate also decreased Palmitoleic acid (PO, C16:1Δ9) content in Peldon (ca. 25% in shoots and 12% in root) and Rothamsted (ca. 39% in shoots and 27% in roots). This, together with the reduction in OA, could also suggest that tri-allate influences the plastid-located steroyl-CoA delta-9 desaturase (SAD) (Figure 1). Looking closely at the content in palmitic acid (PA, C16:0) and stearic acid (SA, C18:0) revealed a differential effect in the two biotypes tested. In Peldon, the content of these two saturated fatty acids is not greatly affected by tri-allate treatment in either shoots or roots, which is consistent with the specific inhibition of fatty acid desaturation hypothesised above. In Rothamsted, PA and SA content are reduced in both shoots and roots (Figure 5), suggesting the inhibition of fatty acid synthesis upstream in the prokaryotic part of the biosynthetic pathway (Figure 1). This would explain the greater reduction in total fatty acid content observed in this ecotype in both shoots and roots after treatment with tri-allate. In Rothamsted, this reduction in C16 and C18 saturated FAs, as well as the general reduction in total FAs in both the shoots and the roots, indicate that tri-allate may act upstream of C16:0-ACP, including on either the plastid-located FAS complex and/or enzymes above (Figure 1). In Peldon treated with tri-allate FAMEs profiles rather suggest the inhibition of the plastidial delta-9 steroyl-ACP desaturase (Figure 1), resulting in the decreased content in C18:1, impacting the profile of all desaturated C18 FAs in shoot and root tissue (Figure 5). The differential effects of tri-allate on VLCFA profiles between both biotypes in shoot and root tissues are interesting but more difficult to explain.

The three herbicides tested in this study had a quantitative effect on surface wax load, but only the two thiocarbamates influenced surface wax composition and cause a qualitative effect (Table 1). This was characterised by an increase in the proportion of FAOHs and decreases in alkanes and aldehydes only visible after treatment with Tri-allate and EPTC, but not FFT. Both aldehydes and alkanes are formed by the decarbonylation (alkane) pathway, whereas fatty alcohols are synthesised by the reducing (alcohol) pathway. Interestingly, the three herbicides tested affected surface wax molecular species with carbon chain lengths of 26 carbons and over, while C22 and C24 compounds were not greatly affected (Table 2). This suggests that another α-oxyacetamide- and thiocarbamate-insensitive elongation system may be responsible for synthesising acyl chains up to 24 carbons when KCS-mediated fatty acid elongation is inhibited. The ELO genes identified in plants (e.g. 4 genes in *A. thaliana* by Nagano et al. (2019)) may be responsible for this activity, but further research to identify and functionally validate this hypothesis *in planta*, specifically in blackgrass, would be required to make this conclusion.

## Materials and Methods

### Alopecurus myosuroides biotypes

The processes used to generate the “purified populations” are fully described in Mellado-Sánchez et al. (2020), Comont et al. (2022), Cai et al. (2023) and Fu et al. (2023). In short, the seed lines were selected specifically to robustly exhibit “Herbicide Sensitive” or “NTSR-only Resistance” phenotypes. The resistant population (Peldon) were derived from individuals originally collected by (Moss, 1990), but selected to exhibit strong NTSR herbicide resistance against acetyl-coenzyme A carboxylase (ACCase)-inhibiting herbicide (fenoxaprop) while carrying the wild-type alleles of all known TSR mutations for acetolactate synthase (ALS) or ACCase. The herbicide-naïve population (Rothamsted) were derived from clones of plants confirmed to be completely sensitive to a panel of herbicides from individuals originally collected from the herbicide-free section of Broadbalk (Moss et al., 2004).

### Seed Sterilisation and Media Preparation

Surface sterilisation of blackgrass seed was performed following an adapted version of adapted from Speakman and Krüger (1983). Seeds were soaked in Terramycin (10 ppm) for 20 hours on a rotator in the dark, rinsed, and then treated with AgNO3 (0.1%) and NaCl (0.5%) solutions before final rinses with sterile water.

For media preparation, Hoagland’s No. 2 Basal Salt Mixture (H2395-10L, Merck) was mixed with MQ water to create a 0.85X solution, and the pH was adjusted to 7.0 with 1N KOH. Agar was weighed and added to bottles for a 0.7% solution. This mixture was then autoclaved and kept at 55 °C. Herbicides were added to each bottle containing 1 L Hoagland’s according to the doses required, starting from the lowest concentration. The herbicide formulations tested in this study included Avadex 480 (480g/l tri-allate), Eptam (800g/l EPTC), and Sunfire (500g/l flufenacet). To each labelled sterile container (E1674.0001 DUCHEFA OS140, Melford), 140 mL media was added and then allowed to cool and solidify before adding sterilised seed.

### Plant growth conditions and sampling procedure

For testing responses to different herbicide doses, 10-12 sterilised seeds were spaced out on the prepared agar media and grown under controlled conditions (17 °C/11 °C with 16 hrs light) for two weeks. For each herbicide dose, there were three replica containers. After two weeks, each seedling was removed gently with forceps from each container and laid between two sheets of acetate; a row from the same container represented one replica. Scanned using a flatbed scanner for image analysis in ImageJ to measure the length of each plant’s root and shoot separately.

For the material used in lipid analysis, referred to as single-dose experiments, an increased number of 18-20 seeds were grown under the same controlled conditions. For each herbicide, three replica control and herbicide-treated containers were used to measure the response as expected. The materials used for analysis came from additional containers; each seedling was carefully removed from the container, and then, using sterilised scissors, roots and shoots were separated from the seed and placed into a pre-weighed glass tube. One tube contained root or shoot material from one replicate. After adding the material, the tubes were reweighed to calculate the fresh weight, then freeze-dried overnight and sealed to store.

### Herbicide dose response testing and analysis

ImageJ was used to quantify root and shoot lengths from scanned images that included a reference ruler. The data was analysed in Excel to calculate growth means, standard errors, t-tests comparing treated and control plants, and comparing the two biotypes. For each treatment replicate, the measured length was divided by the average length of the corresponding control group replicates from the same experimental batch. This normalisation step expressed the data as a ratio relative to the control mean, centring the control values at 1.0. Therefore, treatment values greater than 1.0 indicate increased growth compared to control, while values less than 1.0 represent growth inhibition relative to the untreated state.

The response data were visualised in Excel, and the normalised data were used to identify the herbicide and dose, resulting in ∼40% growth inhibition. This approximate effective dose was chosen as the target dose for herbicide treatments to ensure sufficient plant tissue, especially root tissue, would be available for subsequent analyses while still inducing an herbicidal effect on plant growth.

### Wax extraction, GC-FID/MS and analysis

Gas chromatographic (GC) analysis of fatty acid methyl esters was chosen to rapidly determine oil composition and abundance. The full process of fatty acid methyl ester (FAME) analysis consists of the hydrolysis of lipids, the transesterification of the released fatty acids, injection, separation, identification, and quantitation of the FAMEs. Lipids are extracted and directly esterified in the methylation mixture. Esterification of lipids can be carried out with several reagents based on acid-catalysed or base-catalysed reactions. FAMEs are identified by comparison of their retention times with those of individual purified standards and also by their absolute mass and fractionation pattern in GC coupled with mass-spectrometry. Relative retention times and equivalent chain-length values provide useful information for identification. FAMEs are then quantitated by peak area after correction using individual response factors. Absolute concentrations are determined by adding an internal standard e.g., C17:0 (carbons:desaturations). Compositional changes in fatty acids can reflect the influence of genotype (fatty acid synthesis & lipid metabolism) and environment.

Freeze-dried shoot and root samples were immersed for 60 seconds in 10 mL of chloroform containing 10 mM of docosane (C22 alkane) and eicosanol (C20 primary fatty alcohol) as Internal Standards (IS) to extract surface lipids. Chloroform extracts were transferred to a clean glass tube and evaporated under nitrogen. Samples were derivatised in 100 µL of N,O-bis(trimethylsilyl)trifluoroacetamide): trimethylchlorosilane (99:1; BSTFA/TMS) at 85 °C for 1 hour. Surplus BSTFA-TMCS was evaporated under N2 gas, and the samples dissolved in 200 µL of hexane and transferred to a glass vial for analysis by GC-FID and GC-MS. 1 mL of silylated samples was analysed by GC using a 30 m, 0.25-mm, 0.25 mm HP-1MS capillary column with helium as the carrier gas using the following method: spitless injection, inlet and detector temperatures were set at 325 °C, constant flow rate of 1.5 ml/min, the oven start temperature was set at 50 °C and held for 1 min then increased to 325 °C at a rate of 7 °C/min and the final temperature was held at 325 °C for 15 min. Compounds are identified using a Mass Spectrometer and quantified using a flame-ionization detector (FID). Quantification is based on flame ionization detector peak areas and an internal molecular standard. The total amount of cuticular wax is expressed per unit of fresh weight (FW).

### Lipid extraction

Lipids were extracted from an average of 39.4 mg of leaf or an average of 35.6 mg root material (for sample weights see relevant extraction data file). Glassware was used throughout the procedure. The samples and 1 ml of isopropanol were incubated at 75 °C for 20 minutes. Subsequently, 2 mL of chloroform/methanol (2:1) and 0.7 mL of water were added. After 30 seconds of vortexing, an additional 2 mL of chloroform/water (1:1) was added. The mixture was centrifuged for 3 minutes at 500 g, and the lower chloroform phase was gently transferred to a new tube. Lipids were extracted by adding 1 mL of chloroform to the first tube, repeating the centrifugation, and recovering the lower phase. Both extractions were combined. The chloroform was evaporated using N2 gas while keeping the sample in a 37 °C block. Once all the solvent was evaporated, the lipids were resuspended in 200 μL of chloroform and stored at - 80 °C for subsequent fatty acid analysis.

### Fatty Acid Analysis by GC-FID

Methyl ester derivatives of total fatty acids (FAMEs) were analysed by Gas Chromatography (GC) (Agilent 7890A, Agilent Technologies) using an Agilent J&W 122-2332 column (30 m × 250 µm × 0.25 µm, Agilent Technologies). Inlet and detector temperatures were set to 250 °C, and 1 µL of each sample was analysed using a 15:1 split ratio injection and a constant flow rate of 1.5 mL/minute. The oven temperature cycle has a start temperature of 150 °C held for 2 minutes to allow vaporised samples and the solvent (hexane) to condense at the front of the column. The oven temperature was then increased to 240 °C at 10 °C/minute. The final temperature of 240 °C was held for one minute and 50 s, giving a total run time of 12 min and 50 s per sample. FAMEs were detected using a Flame Ionisation Detector (FID). Chromatograms were analysed using the offline session of the Agilent ChemStation software (Agilent Technologies, USA). The peak area of each FAME was normalized to the internal standard and further normalized to the weight of the initial sample. The retention time and identity of each peak were calibrated using the Supelco® 37 Component FAME Mix (certified reference material TraceCERT®, Sigma-Aldrich®).

### Fatty Acid Analysis by GC-MS

The samples were injected on an Agilent 7890A gas chromatograph coupled to an Agilent 5975C Mass Spectrometer on the same column as wax analyses with the following difference: inlet and detector temperatures were set at 290 °C, constant flow rate of 1.5 ml/min, the oven start temperature was set at 50 °C and held for 1 min then increased to 290 °C at a rate of 10 °C/min and the final temperature was held at 290 °C for 2min. VLCFAs were detected using selective ion monitoring methods and quantified using the most abundant ion in the mass spectra i.e. the m/z = 74 corresponding to McLafferty rearrangement ion for saturated compounds (C20:0-C32:0) or the m/z = 57 ion for monounsaturated compounds (C20:1-C32:1).

## Acknowledgments and Funding

Rothamsted Research receives strategic funding from the Biotechnology and Biological Sciences Research Council of the United Kingdom (BBSRC). We acknowledge support from the Growing Health Institute Strategic Programme [BB/X010953/1; BBS/E/RH/230003A]. This research was supported by Gowan Crop Protection Limited, where authors BH and LC are currently employed. The company produces tri-allate and EPTC. The authors would like to acknowledge Christian Harrison and Margaret (Peggy) McGroary for their help in sampling the plant material from the discriminatory dose experiments as well as Sumit Sethi who helped with the wax and FAMEs extractions.

## Author contributions

DM, FB, LC, and BH conceived the project and designed the study. HB and DM adapted the agar-based system for use with pre-emergent herbicides, grew the plants in this system, and sampled them. DM and HB analysed the physiological changes that resulted from the different herbicide dose assays and chose the discriminatory doses. HB and FB extracted waxes and lipids for and performed the analysis by GC-FID/MS. FB, RH planned and supervised data analysis and production for wax and lipid measurements. DM was the overall project manager. HB, DM, FB and RH wrote the initial manuscript draft and LC and BH contributed editing and improvements. All authors read and approved the final manuscript and agree to be accountable for all aspects of the work in ensuring that questions related to the accuracy or integrity of any part of the work are appropriately investigated and resolved.

## Data availability

The data supporting the findings of this study are available in the article and its supplementary information files.

## Dive Curated Terms

The following phenotypic, genotypic, and functional terms are of significance to the work described in this paper:

- Flufenacet CHEBI: CHEBI:81920 - https://www.ebi.ac.uk/chebi/searchId.do;?chebiId=CHEBI:81920
- EPTC CHEBI: CHEBI:4738 - https://www.ebi.ac.uk/chebi/searchId.do?chebiId=CHEBI:4738
- Tri-allate CHEBI: CHEBI:81978 - https://www.ebi.ac.uk/chebi/searchId.do?chebiId=CHEBI:81978
- KEGG Pathway Fatty Acid Metabolism – Reference Pathway - https://www.genome.jp/pathway/map01212 and https://www.genome.jp/pathway/bdi00061
- Ketoacyl-ACP Synthase III (KASIII) - https://www.genome.jp/entry/prel:PRELSG_0408700
- Ketoacyl-ACP Reductase (KAR), https://www.arabidopsis.org/locus?key=28873
- Hydroxyacyl-ACP Dehydrase (HAD), https://www.genome.jp/dbget-bin/www_bget?ec:4.2.1.59
- Enoyl-ACP Reductase (ER), https://www.arabidopsis.org/locus?key=35306
- Ketoacyl-ACP Synthase I (KASI), https://www.arabidopsis.org/locus?key=134125
- Ketoacyl-ACP Synthase II (KASII), https://www.ncbi.nlm.nih.gov/protein/AAL91174.1/
- Stearoyl-ACP Desaturase (SAD), https://www.arabidopsis.org/locus?key=29774
- Acyl-ACP Thioesterase B (FATB), https://www.arabidopsis.org/locus?key=137248
- Acyl-ACP Thioesterase A (FATA), https://www.arabidopsis.org/locus?key=38410
- Fatty Acid Export (FAX1), https://www.arabidopsis.org/locus?key=37036
- Long-Chain Acyl-CoA Synthetase (LACS9), https://www.arabidopsis.org/locus?key=137629
- Oleate Desaturase (FAD2), https://www.arabidopsis.org/locus?key=39962
- Linoleate Desaturase (FAD3), https://www.arabidopsis.org/locus?key=26541

## Author notes

Dana MacGregor corresponding author can be contacted by email at.

The author responsible for distribution of materials integral to the findings presented in this article in accordance with the policy described in the Instructions for Authors (https://academic.oup.com/plphys/pages/General-Instructions) is: Dana MacGregor.

Two authors work for Gowan Crop Protection Limited, a company which produces the herbicides tri-allate (Avadex) and EPTC (Eptam).

## Abbreviations

“Roth”: Blackgrass biotype “Rothamsted” – Herbicide-naïve population described in Mellado-Sánchez et al. (2020) and Fu et al. (2023). In short, the Roth purified population was derived from seeds collected from the herbicide-free section of Broadbalk field trials (Moss et al., 2004) where plants were clonally propagated and the clones of plants that did not survive a panel of herbicides were allowed to bulk cross in isolation.

“PELD”: Blackgrass biotype “Peldon” – Non-target site resistant population described in Mellado-Sánchez et al. (2020) and Fu et al. (2023). In short, this population is derived from seeds collected from Moss (1990) from location “Peldon” where plants were clonally propagated and the clones of plants that exhibited strong NTSR herbicide resistance but did not carry any known TSR mutations were allowed to bulk cross in isolation.

FW: Fresh Weight
IS: Internal standard min: Minute
GC-FID: Gas Chromatography coupled to a Flame Ionisation Detector
GC-MS: Gas Chromatography coupled to a Mass Spectrometry Detector
VLC-FAOH: Very Long Chain Fatty Alcohols
VLCFA: Very Long Chain Fatty Acid
FFT: Flufenacet / α-oxyacetamide
EPTC: S-ethyl dipropylthiocarbamate
Tri-allate: S-(2,3,3-trichloroprop-2-en-1-yl) di(propan-2-yl) carbamothioate

## Supplementary Figures

**Supplementary Figure 1:**
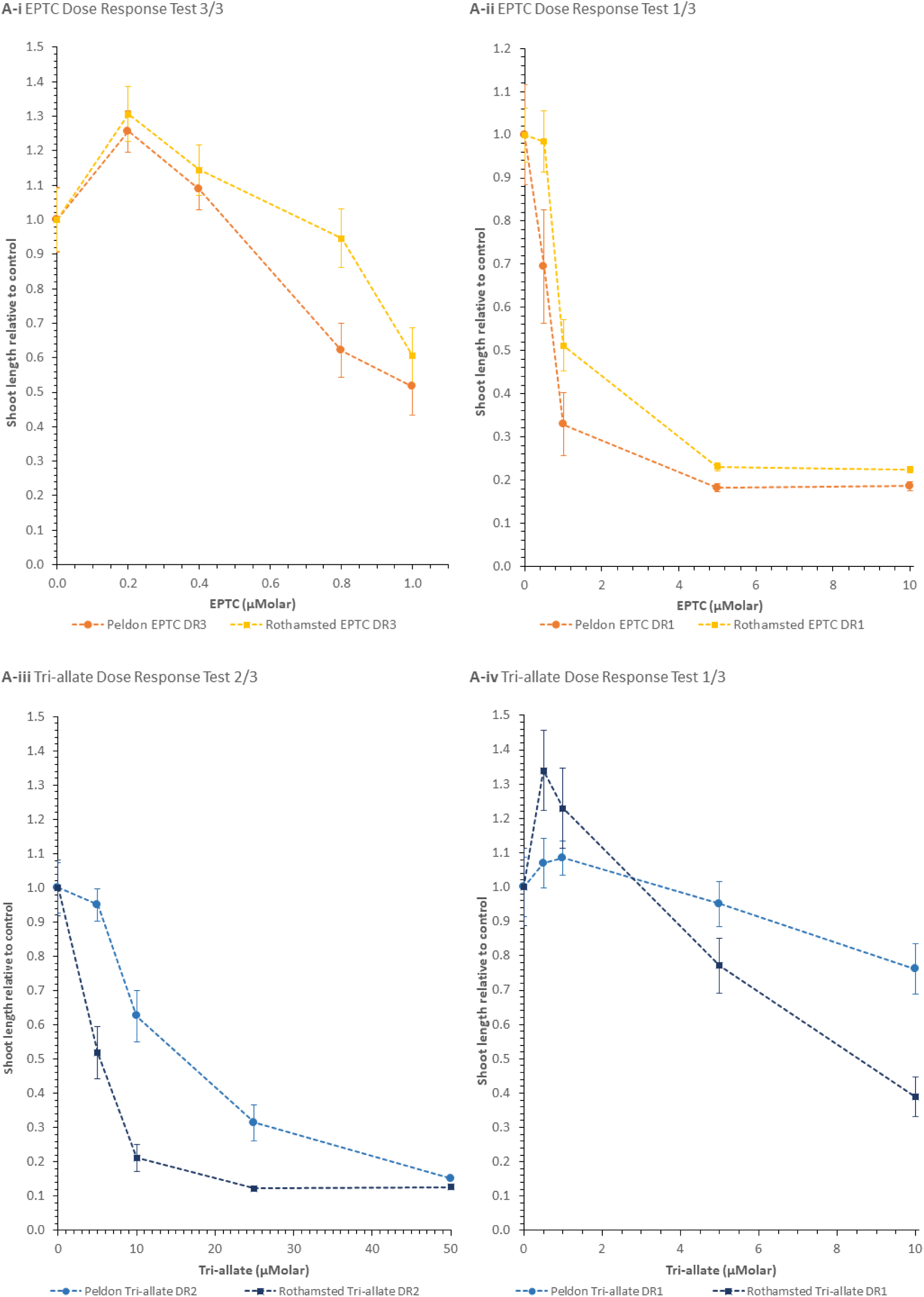
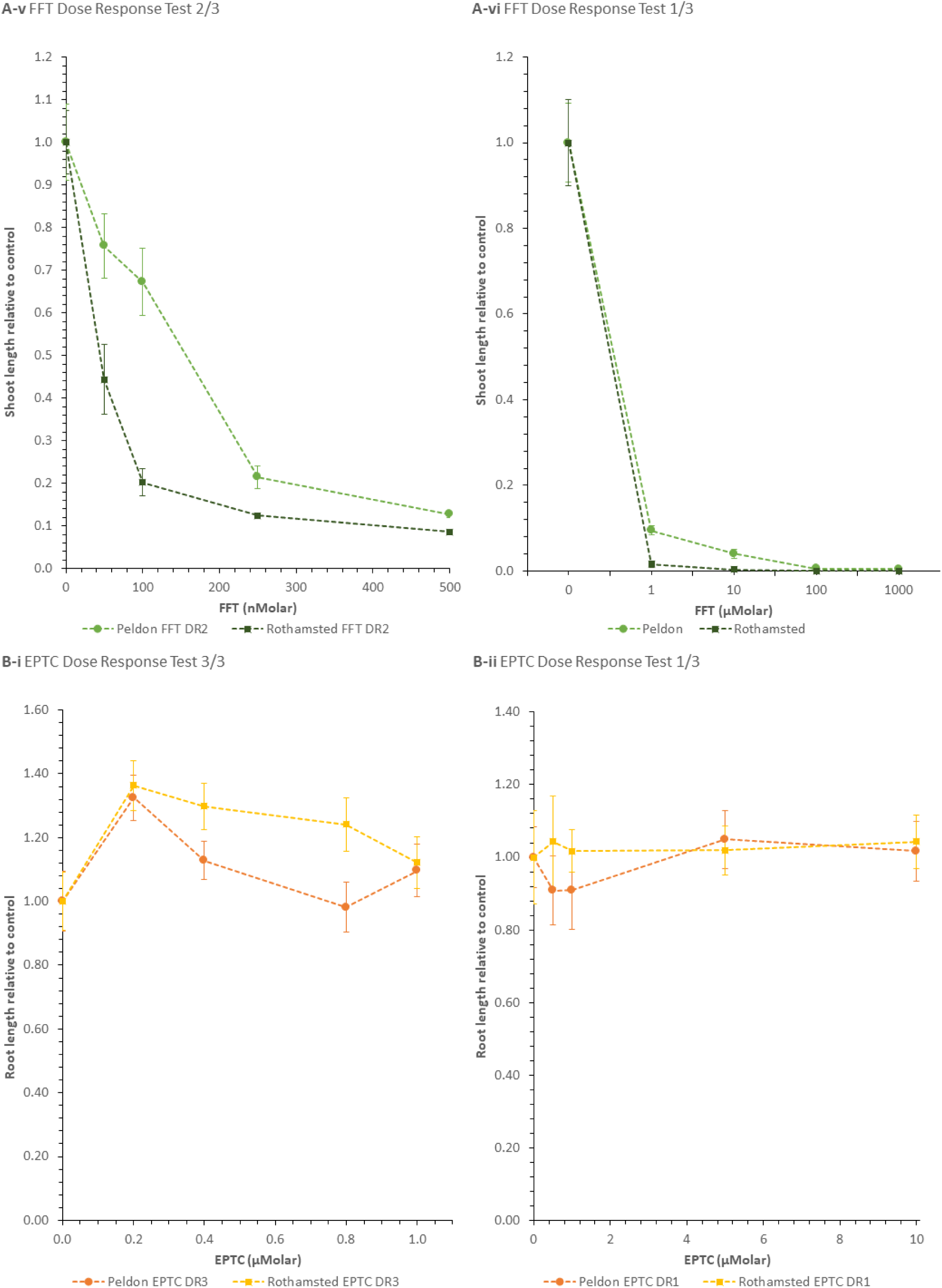
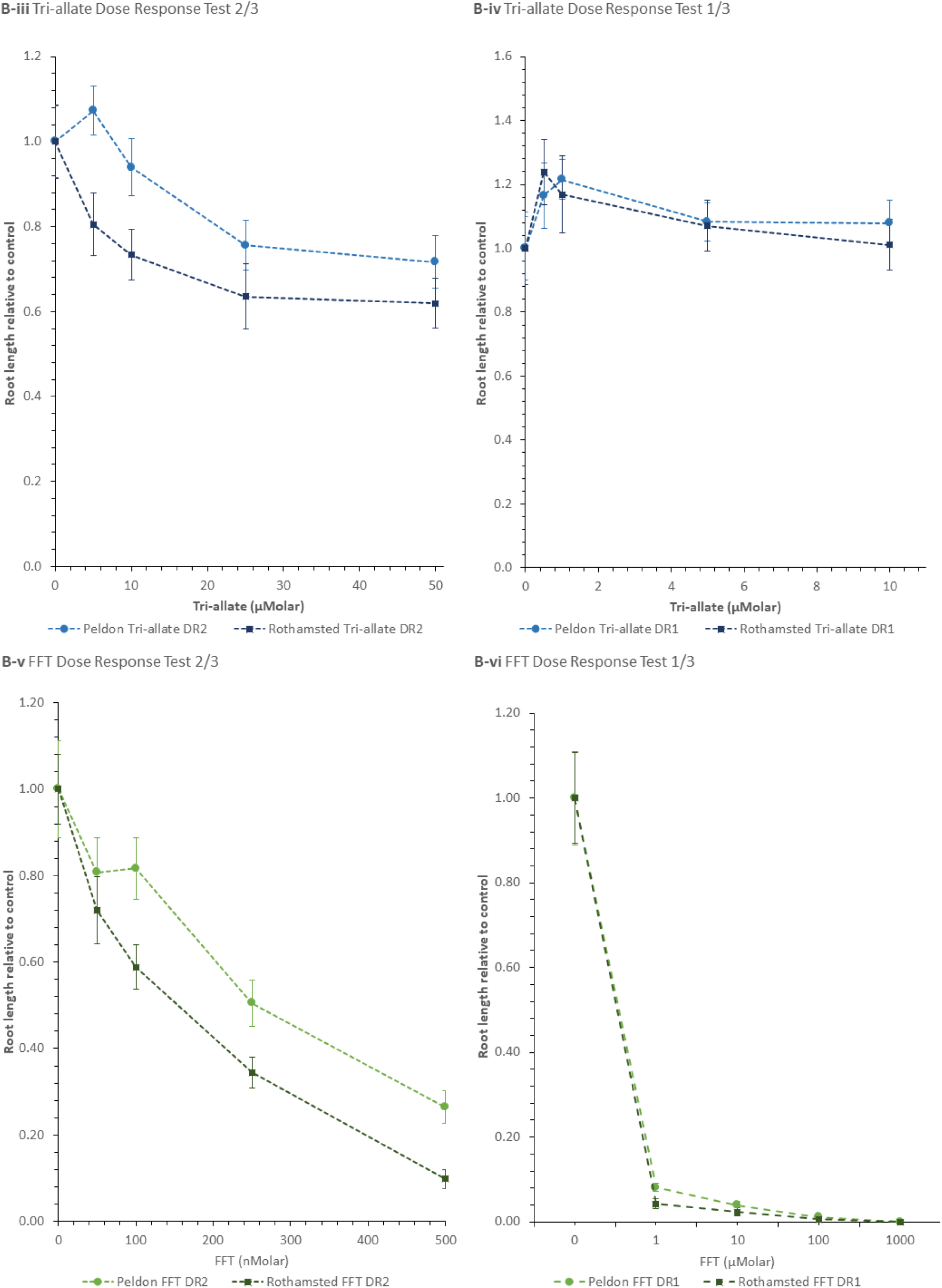
Additional dose testing curves showing the different effects of the pre-emergent herbicides on the two blackgrass biotypes on (A) shoots and (B) roots. Showing average lengths relative to the biotype controls ± standard error for dose-response tests in EPTC (i/ii), Tri-allate (iii/iv) and FFT (v/vi).

**Supplementary Figure 2:**
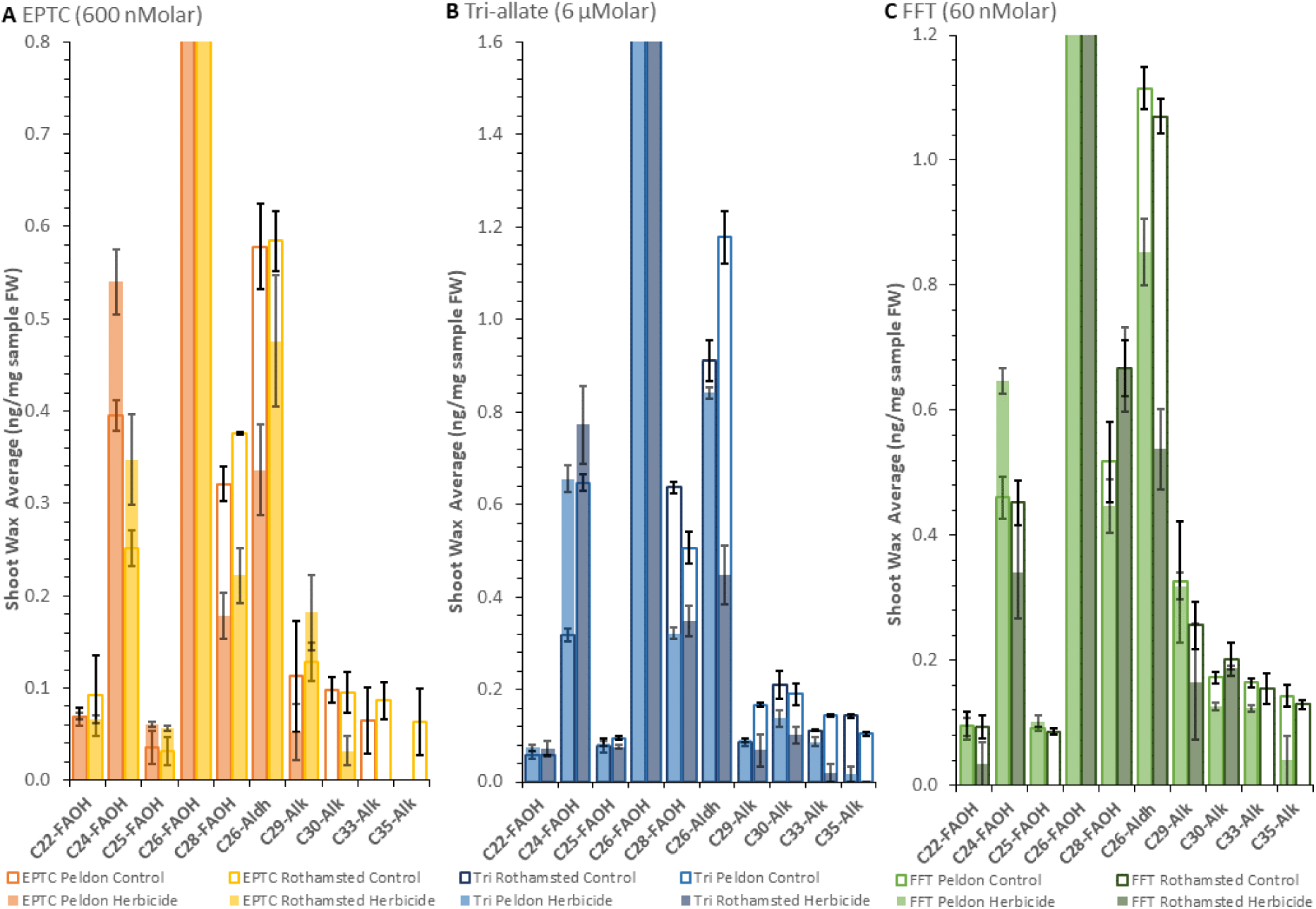
Shoot wax low abundance species, Rothamsted and Peldon average composition ng/mg FW sample with and without herbicide treatment with (A) EPTC, (B) Tri-allate and (C) FFT. Analysis of data generated through GC-FID. The outlines represent the control, while the fill represents the response to herbicide treatment n=3.

**Supplementary Figure 3:**
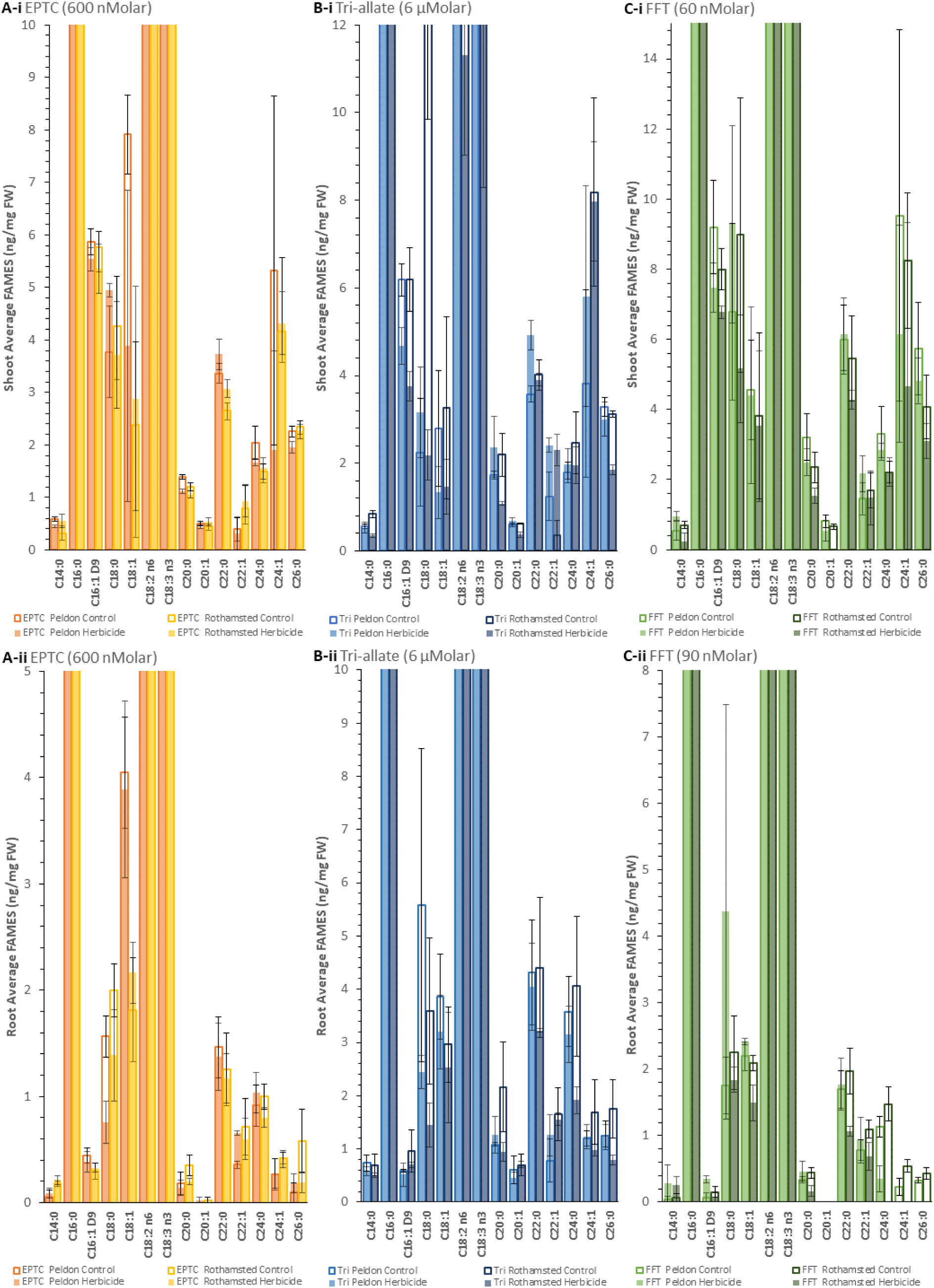
Lower abundance species blackgrass FAMES (i)shoot and (ii) root Following Herbicide Treatment (A) EPTC, (B) Tri-allate (Tri) and (C) FFT. Analysed using GC-FID. Control samples not treated with herbicide are included for baseline comparison. Each data point represents the mean of three biological replicates, except for in (A-i) Peldon Herbicide treated and (B-ii) Tri-allate Control, where n=2, with error bars indicating a standard error.

## Supplementary Tables

**Supplementary Table 1:**
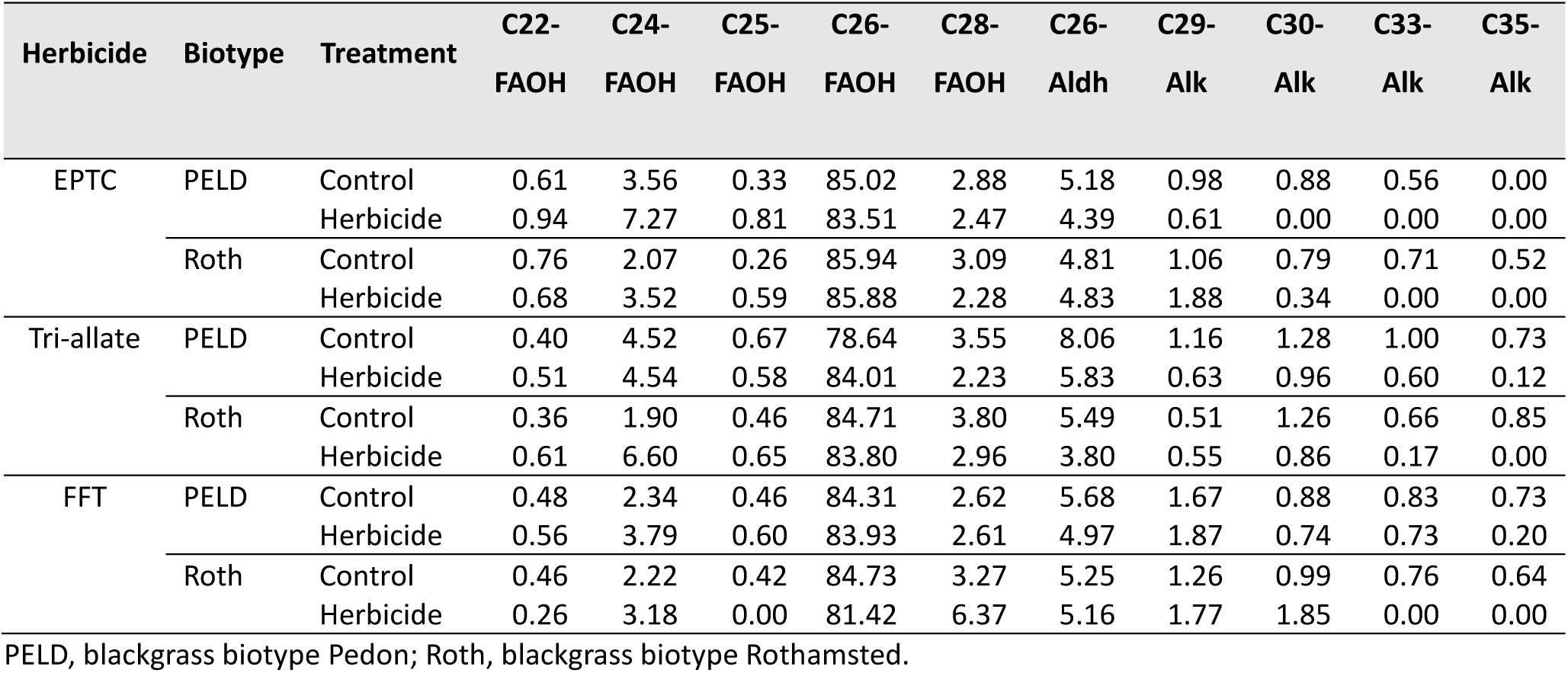
Percentage of total wax species in blackgrass shoots with and without herbicide treatment.

| Herbicide | Biotype | Treatment | C22-FAOH | C24-FAOH | C25-FAOH | C26-FAOH | C28-FAOH | C26-Aldh | C29-Alk | C30-Alk | C33-Alk | C35-Alk |
| --- | --- | --- | --- | --- | --- | --- | --- | --- | --- | --- | --- | --- |
| EPTC | PELD | Control | 0.61 | 3.56 | 0.33 | 85.02 | 2.88 | 5.18 | 0.98 | 0.88 | 0.56 | 0.00 |
|  |  | Herbicide | 0.94 | 7.27 | 0.81 | 83.51 | 2.47 | 4.39 | 0.61 | 0.00 | 0.00 | 0.00 |
|  | Roth | Control | 0.76 | 2.07 | 0.26 | 85.94 | 3.09 | 4.81 | 1.06 | 0.79 | 0.71 | 0.52 |
|  |  | Herbicide | 0.68 | 3.52 | 0.59 | 85.88 | 2.28 | 4.83 | 1.88 | 0.34 | 0.00 | 0.00 |
| Tri-allate | PELD | Control | 0.40 | 4.52 | 0.67 | 78.64 | 3.55 | 8.06 | 1.16 | 1.28 | 1.00 | 0.73 |
|  |  | Herbicide | 0.51 | 4.54 | 0.58 | 84.01 | 2.23 | 5.83 | 0.63 | 0.96 | 0.60 | 0.12 |
|  | Roth | Control | 0.36 | 1.90 | 0.46 | 84.71 | 3.80 | 5.49 | 0.51 | 1.26 | 0.66 | 0.85 |
|  |  | Herbicide | 0.61 | 6.60 | 0.65 | 83.80 | 2.96 | 3.80 | 0.55 | 0.86 | 0.17 | 0.00 |
| FFT | PELD | Control | 0.48 | 2.34 | 0.46 | 84.31 | 2.62 | 5.68 | 1.67 | 0.88 | 0.83 | 0.73 |
|  |  | Herbicide | 0.56 | 3.79 | 0.60 | 83.93 | 2.61 | 4.97 | 1.87 | 0.74 | 0.73 | 0.20 |
|  | Roth | Control | 0.46 | 2.22 | 0.42 | 84.73 | 3.27 | 5.25 | 1.26 | 0.99 | 0.76 | 0.64 |
|  |  | Herbicide | 0.26 | 3.18 | 0.00 | 81.42 | 6.37 | 5.16 | 1.77 | 1.85 | 0.00 | 0.00 |

PELD, blackgrass biotype Pedon; Roth, blackgrass biotype Rothamsted.

**Supplementary Table 2.1:** mass% FAMES Shoot.

| Herbicide | Biotype | Treatment | C14:0 | C16:0 | C16:1 D9 | C18:0 | C18:1 | C18:2 n6 | C18:3 n3 | C20:0 | C20:1 | C22:0 | C22:1 | C24:0 | C24:1 | C26:0 |
| --- | --- | --- | --- | --- | --- | --- | --- | --- | --- | --- | --- | --- | --- | --- | --- | --- |
| EPTC | PELD | Control | 0.17 | 11.50 | 1.71 | 1.10 | 2.31 | 19.04 | 59.69 | 0.41 | 0.15 | 0.98 | 0.11 | 0.60 | 1.56 | 0.66 |
|  |  | Herbicide | 0.14 | 11.21 | 1.75 | 1.56 | 1.23 | 18.89 | 61.70 | 0.35 | 0.14 | 1.18 | 0.10 | 0.53 | 0.60 | 0.62 |
|  | Roth | Control | 0.10 | 10.51 | 1.91 | 1.41 | 0.79 | 19.31 | 61.63 | 0.40 | 0.15 | 0.88 | 0.27 | 0.49 | 1.38 | 0.78 |
|  |  | Herbicide | 0.19 | 11.60 | 1.86 | 1.29 | 1.00 | 20.48 | 58.80 | 0.37 | 0.19 | 1.07 | 0.33 | 0.54 | 1.50 | 0.78 |
| Tri-allate | PELD | Control | 0.15 | 10.70 | 1.72 | 0.62 | 0.78 | 18.20 | 63.38 | 0.48 | 0.17 | 0.99 | 0.34 | 0.50 | 1.06 | 0.91 |
|  |  | Herbicide | 0.40 | 28.49 | 3.54 | 2.39 | 1.01 | 16.04 | 32.06 | 1.80 | 0.51 | 3.74 | 1.83 | 1.50 | 4.41 | 2.28 |
|  | Roth | Control | 0.25 | 10.91 | 1.81 | 3.66 | 0.96 | 18.94 | 57.34 | 0.64 | 0.18 | 1.17 | 0.10 | 0.72 | 2.39 | 0.91 |
|  |  | Herbicide | 0.43 | 36.93 | 4.62 | 2.69 | 1.81 | 13.93 | 15.66 | 1.33 | 0.46 | 4.81 | 2.83 | 2.41 | 9.81 | 2.27 |
| FFT | PELD | Control | 0.10 | 10.59 | 1.62 | 1.20 | 0.81 | 18.26 | 62.11 | 0.57 | 0.14 | 1.06 | 0.26 | 0.59 | 1.68 | 1.01 |
|  |  | Herbicide | 0.19 | 11.79 | 1.48 | 1.84 | 0.87 | 19.40 | 59.45 | 0.49 | 0.11 | 1.22 | 0.43 | 0.56 | 1.22 | 0.95 |
|  | Roth | Control | 0.16 | 11.08 | 1.81 | 2.03 | 0.87 | 18.86 | 59.60 | 0.53 | 0.15 | 1.24 | 0.39 | 0.50 | 1.87 | 0.92 |
|  |  | Herbicide | 0.06 | 11.60 | 1.66 | 1.27 | 0.86 | 19.98 | 60.34 | 0.38 | 0.00 | 1.05 | 0.36 | 0.55 | 1.14 | 0.76 |

PELD, blackgrass biotype Pedon; Roth, blackgrass biotype Rothamsted.

**Supplementary Table 2.2:** mass% FAMES Root.

| Herbicide | Biotype | Treatment | C14:0 | C16:0 | C16:1 D9 | C18:0 | C18:1 | C18:2 n6 | C18:3 n3 | C20:0 | C20:1 | C22:0 | C22:1 | C24:0 | C24:1 | C26:0 |
| --- | --- | --- | --- | --- | --- | --- | --- | --- | --- | --- | --- | --- | --- | --- | --- | --- |
| EPTC | PELD | Control | 0.09 | 18.82 | 0.69 | 2.42 | 6.26 | 40.55 | 26.11 | 0.28 | 0.00 | 2.26 | 0.55 | 1.41 | 0.42 | 0.14 |
|  |  | Herbicide | 0.16 | 18.82 | 0.68 | 1.35 | 6.92 | 40.01 | 25.49 | 0.25 | 0.04 | 2.45 | 1.16 | 1.84 | 0.49 | 0.32 |
|  | Roth | Control | 0.30 | 20.97 | 0.50 | 3.35 | 3.04 | 42.06 | 22.50 | 0.59 | 0.04 | 2.10 | 1.20 | 1.67 | 0.71 | 0.98 |
|  |  | Herbicide | 0.40 | 20.26 | 0.61 | 2.60 | 4.06 | 42.45 | 23.31 | 0.38 | 0.00 | 2.19 | 1.12 | 1.50 | 0.77 | 0.35 |
| Tri-allate | PELD | Control | 0.47 | 19.70 | 0.38 | 3.57 | 2.48 | 41.19 | 24.01 | 0.69 | 0.39 | 2.76 | 0.50 | 2.30 | 0.77 | 0.80 |
|  |  | Herbicide | 0.43 | 22.02 | 0.38 | 1.80 | 2.35 | 41.63 | 22.09 | 0.93 | 0.33 | 2.98 | 0.93 | 2.32 | 0.91 | 0.91 |
|  | Roth | Control | 0.35 | 21.77 | 0.48 | 1.81 | 1.50 | 39.83 | 25.97 | 1.09 | 0.35 | 2.22 | 0.83 | 2.05 | 0.85 | 0.88 |
|  |  | Herbicide | 0.47 | 25.06 | 0.65 | 1.34 | 2.34 | 42.35 | 18.40 | 0.88 | 0.65 | 2.99 | 1.44 | 1.78 | 0.91 | 0.74 |
| FFT | PELD | Control | 0.09 | 21.27 | 0.15 | 3.52 | 4.39 | 40.04 | 21.53 | 0.68 | 0.00 | 3.39 | 1.56 | 2.26 | 0.46 | 0.66 |
|  |  | Herbicide | 0.47 | 22.17 | 0.59 | 7.45 | 4.12 | 40.79 | 18.41 | 0.79 | 0.00 | 3.02 | 1.59 | 0.60 | 0.00 | 0.00 |
|  | Roth | Control | 0.10 | 22.18 | 0.23 | 3.42 | 3.19 | 42.68 | 19.13 | 0.67 | 0.00 | 3.01 | 1.67 | 2.25 | 0.83 | 0.66 |
|  |  | Herbicide | 0.46 | 20.53 | 0.13 | 3.33 | 2.70 | 49.71 | 19.67 | 0.30 | 0.00 | 1.92 | 1.24 | 0.00 | 0.00 | 0.00 |

PELD, blackgrass biotype Pedon; Roth, blackgrass biotype Rothamsted.

**Supplementary Table 3:** Percentage change in FAMES split into FA and VLCFA ng/mg FW.

| Herbicide | Biotype | % change in ng/mg FW of shoots |  |  | % change in ng/mg FW of roots |  |  |
| --- | --- | --- | --- | --- | --- | --- | --- |
|  |  | FA | VLCFA | Total | FA | VLCFA | Total |
| EPTC | PELD | -6.7% | -27.2% | **-7.6%** | -14.5% | 12.4% | **-13.1%** |
|  | Roth | -5.0% | 4.8% | **-4.6%** | -9.6% | -22.5% | **-10.5%** |
| Tri-allate | PELD | -67.8% | 32.0% | **-63.4%** | -13.9% | -1.3% | **-12.8%** |
|  | Roth | -80.7% | -7.3% | **-76.2%** | -46.2% | -38.3% | **-45.6%** |
| FFT | PELD | -10.5% | -16.3% | **-10.8%** | 21.0% | -22.0% | **17.1%** |
|  | Roth | -6.2% | -30.0% | **-7.5%** | -10.6% | -67.8% | **-15.8%** |

Summed FA (C14:0 to C18:3), VLCFA (C20:0 to C26:0) and total FAMES from values from Tables 3.1 and 3.2, calculated percentage change using the formula: $\frac{((\text{control} - \text{herbicide})/\text{control}) * -1}{1} * 100$ . The multiplication by minus one helps illustrate the direction of change.

T-test results behind Figure 3. (control vs. treated, relative to control)

|  | Treatment | PELD Shoot | PELD Root | Roth Shoot | Roth Root |
| --- | --- | --- | --- | --- | --- |
| Tri-allate | Control | 1.00E+00 | 1.00E+00 | 1.00E+00 | 1.00E+00 |
| | 6 μMolar | 3.57E-01 | 4.43E-01 | **4.75E-05** | 8.71E-01 |
| EPTC | Control | 1.00E+00 | 1.00E+00 | 1.00E+00 | 1.00E+00 |
|  | 600 nMolar | **1.18E-03** | 2.20E-01 | 1.78E-01 | 7.98E-01 |
| FFT 60 | Control | 1.00E+00 | 1.00E+00 | 1.00E+00 | 1.00E+00 |
|  | 60 nMolar | **8.46E-04** | **2.85E-02** | **1.27E-10** | 6.84E-01 |
| FFT 90 | Control | 1.00E+00 | 1.00E+00 | 1.00E+00 | 1.00E+00 |
|  | 90 nMolar | **2.48E-04** | 5.48E-02 | **5.46E-09** | **1.36E-03** |

PELD, blackgrass biotype Pedon; Roth, blackgrass biotype Rothamsted. Bold font, $p < 0.05$ .

Additional between-biotype t-tests from the single-dose experiments for both the raw length data and the relative to control, comparing blackgrass biotypes Peldon (PELD) vs. Rothamsted (Roth).

|  | Treatment | Shoot PELD vs. Roth t-test result | Relative to control shoot PELD vs. Roth t-test result |
| --- | --- | --- | --- |
| EPTC | Control | 2.72E-01 | 1.00E+00 |
|  | 600 nMolar | **2.78E-02** | 1.61E-01 |
| Tri-allate | Control | **2.15E-04** | 1.00E+00 |
| | 6 μMolar | 4.48E-01 | **3.98E-03** |
| FFT 60 | Control | 8.67E-02 | 1.00E+00 |
|  | 60 nMolar | **1.07E-02** | **1.14E-03** |
| FFT 90 | Control | 1.98E-01 | 1.00E+00 |
|  | 90 nMolar | **1.53E-03** | **5.66E-04** |
|  | Treatment | Root PELD vs. Roth t-test result | Relative to control root PELD vs. Roth t-test result |
| EPTC | Control | 2.71E-01 | 1.00E+00 |
|  | 600 nMolar | 5.76E-02 | 3.55E-01 |
| Tri-allate | Control | **4.75E-03** | 1.00E+00 |
| | 6 μMolar | **4.55E-02** | 5.88E-01 |
| FFT 60 | Control | 7.00E-01 | 1.00E+00 |
|  | 60 nMolar | **3.01E-02** | **1.10E-02** |
| FFT 90 | Control | 5.63E-01 | 1.00E+00 |
|  | 90 nMolar | 7.73E-02 | **1.99E-02** |

